# Immunity and bacterial recruitment in plant leaves are parallel processes that together shape sensitivity to temperature stress

**DOI:** 10.1101/2024.06.10.598336

**Authors:** Jisna Jose, Erik Teutloff, Teresa Mayer, Simrat Naseem, Emanuel Barth, Rayko Halitschke, Manja Marz, Matthew T. Agler

**Author notes:** Corresponding author: Matthew T. Agler The Plant Microbiosis Lab, Institute of Microbiology Friedrich Schiller University Jena Neugasse 23 07743 Jena, Germany.

## Abstract

Rising global temperatures necessitate developing resilient crops with better adaptability to changing climates. Under elevated temperatures, plant immunity is downregulated, increasing risk of foliar pathogen attack. Manipulating plant defense hormones is one way to mitigate this detrimental effect. However, it is unclear how plant immunity interacts with plant microbiome assembly and how temperature will thus affect overall plant health and stability. In this study, we compared two *Arabidopsis thaliana* genotypes that feature divergent strategies for recruitment of commensal bacteria from natural soil. NG2, an *A. thaliana* ecotype we collected from Jena, Germany, was grown in its native soil and compared to CLLF, a genotype that recruits higher bacterial loads and higher bacterial diversity but without any dysbiotic phenotype. CLLF hyperaccumulates salicylic acid (SA) and jasmonates, has constitutively upregulated innate defenses, and shows increased resistance to necrotrophic fungal and hemi-biotrophic bacterial pathogens, indicating that pathogen immunity and non-pathogen recruitment function in parallel. Some of its leaf bacteria can utlize SA as a carbon source, suggesting that immunity and recruitment may even be linked by chemical hormones. CLLF exhibits high tolerance to heat stress in comparison to the NG2, with SA-associated defense processes remaining active under heat. Synthetic community (SynCom) experiments revealed that when the taxonomic diversity of bacteria available to CLLF is artificially reduced, resilience to heat stress is compromised, leading to dysbiosis. However, this dysbiosis does not occur in CLLF with a full SynCom or in the NG2 with any SynCom. These findings suggest that the downregulation of defenses in response to heat may contribute to the avoidance of dysbiosis caused by certain leaf bacteria, while full bacteriome taxonomic diversity can help maintain balance.

## Background

Extreme conditions such as high temperature, drought, salinity, and high humidity due to climate change may adversely affect agriculture and crop productivity. One reason is that changes in ambient growth conditions that affect plant physiology and development, in turn, can alter plant-microbe interactions, an inter-dependence known as the “disease triangle” (1). Altered plant-microbe interactions carry the risk that balance in the microbiome may be lost, leading to dysbiosis, including devastating plant diseases and loss of productivity. Thus, it is important to understand how plants establish and maintain balance in the microbiome and how environmental factors will affect that balance.

Leaf-colonizing bacteria are an important component of the microbiome with regard to plant health because they protect plants but also must be regulated to prevent disease (2). Plants use a variety of mechanisms to recruit bacteria and maintain balance. The phyllosphere (leaf) bacteriome is assembled from both soil-born and air-born inocula in either a deterministic or stochastic manner (3, 4). Plant defenses, besides establishing immunity to pathogens, also plays critical roles in maintaining balance in the bacteriome. For one, the interaction of commensal and opportunistic pathogenic bacteria with immune components like MAMP-triggered immunity helps prevent dysbiosis by preventing overgrowth of opportunistic pathogens (5–7). Additionally, secondary metabolites can contribute to assembly via their toxicity (8) and likely by serving as resources for bacteria particularly adapted to use them (9, 10). However, how plant immunity against (opportunistic) pathogens is more broadly linked to the process of bacteriome assembly in healthy leaves remains unclear.

The plant immune system is itself affected by the abiotic environment. Although the mechanisms linking abiotic stressors to plant immunity are still underexplored, the effect of temperature is perhaps best studied. For example, at moderately high temperatures (30°C vs. 23°C), the production of salicylic acid (SA), a major plant defense hormone, is suppressed (11). Jasmonic acid (JA) biosynthesis is also involved in temperature-dependent susceptibility of some diseases (12). Abscisic acid (ABA) biosynthesis was not affected by heat stress, but ABA does negatively affect R gene-mediated immune responses at high temperatures (13). Because of compromised immunity, these changes can lead to increased susceptibility to pathogens. Some microbes help plants mitigate heat stress by making them more resilient (14, 15), suggesting that the leaf microbiome could offset some adverse effects of increased temperature.

We hypothesized that the balance established by plant immune components during the natural recruitment of leaf bacteriomes plays a crucial role in resilience to temperature stress. While the composition of plant-associated microbiomes is known to influence host health, the specific role of naturally recruited soil microbiomes in maintaining balance and preventing dysbiosis remains poorly explored. To investigate this, we first studied how plant physiology shapes microbiome balance by analyzing CLLF, a genetically distinct *A. thaliana* genotype that naturally recruits an alternative leaf bacteriome. We then examined its immune signaling, pathogen resistance, and response to temperature stress. Finally, we tested how microbiome composition, particularly the diversity of bacteria recruited from natural soil, contributes to maintaining balance and compensating for altered immune responses under heat stress.

## Results

### An unusual A. thaliana Genotype Displays an Altered Leaf Bacteriome

We compared whole leaf bacteriome assembly in the previously described *A. thaliana* genotype NG2 (10) and the unusual genotype CLLF in the natural NG2 soil (collected near its original isolation site). We used a plastic barrier to separate rosettes from soil and to ensure bacterial colonization is not due to increased direct leaf-soil contact (see methods). In 3-week-old plants, bacterial loads in CLLF leaves were approximately 100-fold higher compared to NG2 according to both CFU counts and hamPCR (16) (Fig. 1B and 1C, p<0.01 and p<0.05, respectively). In addition, CLLF recruited a more diverse set of bacterial genera than NG2 (Fig. 1D, Chao1 p < 0.01) while maintaining a similar evenness (Fig. 1D, Shannon and Simpson indices). Furthermore, CLLF leaf bacteriome samples were more densely clustered and distinctly separated from NG2 in a canonical correspondence analysis (CCA) of Bray-Curtis distances (Fig. 1E, 15.4% variation explained by genotype, PERMANOVA, p = 0.008, 999 permutations). The distinct beta diversity patterns, higher alpha diversity, and increased bacterial loads together confirm that under identical conditions, CLLF recruits an altered leaf microbiota. Similar results were found in three independent experiments conducted without the plastic barrier (Supplementary Fig. 2).

**Fig. 1.**
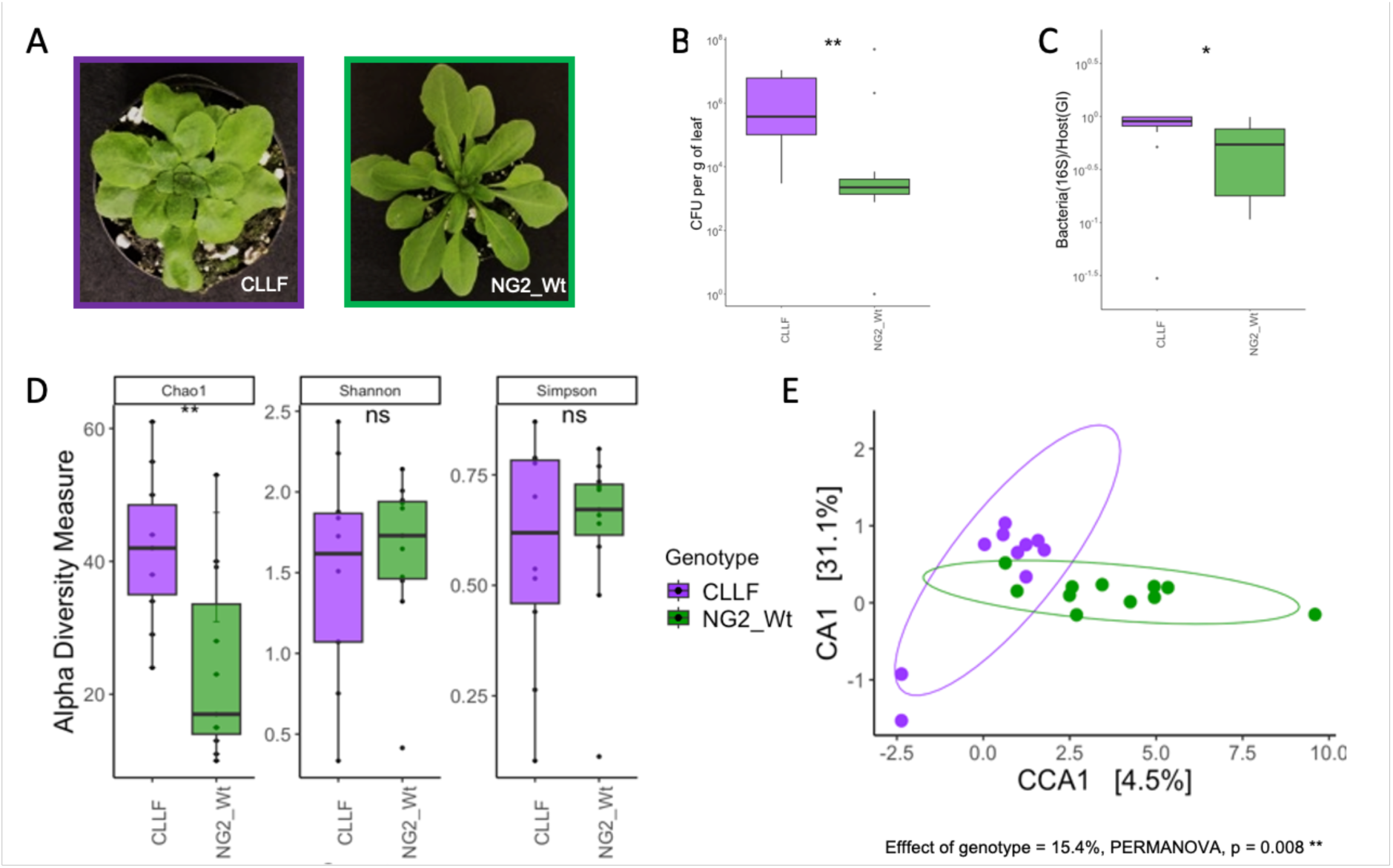
A genetically distinct A. thaliana genotype (CLLF) recruits a distinctive leaf bacteriome. A) Representative images showing the phenotype of CLLF, a genotype with curled and compressed leaves and late flowering. B) Bacterial loads of CLLF and NG2 were measured as colony-forming units (CFU) per gram of fresh leaf weight (n = 12). C) Bacteria-to-host ratio of CLLF and NG2, calculated using hamPCR amplicon sequencing data, representing the fraction of total bacterial 16S rRNA gene reads relative to the sum of bacterial 16S rRNA gene and *A. thaliana* GI (GIGANTEA) reads per sample (n = 12). D) Alpha diversity measures of the CLLF and NG2 leaf bacteriome, based on 16S rRNA gene amplicon sequencing data grouped at the genus level. E) Canonical correspondence analysis (CCA) plot based on Bray-Curtis distances between samples constrained for genotype, generated using 16S rRNA gene amplicon sequencing data grouped at the genus level. 15.4% of variance is explained by genotype (PERMANOVA with 999 permutations), p-value < 0.01. (CLLF; purple, NG2; green, ns = not significant, * p<0.05, ** p<0.01, *** p<0.001).

### CLLF’s altered recruitment is not dysbiotic

To determine whether the high bacterial loads but healthy CLLF phenotype indicated a more taxonomically balanced bacteriome—rather than the over-colonization seen in previously described “dysbiotic” genotypes (characterized by the dominance of a few opportunistic pathogens)—we examined the taxonomic composition of the CLLF leaf bacteriome. Overall, CLLF recruited bacterial families similar to NG2 (Fig. 2A). However, multiple families were significantly more abundant in CLLF (p ≤ 0.01) (Fig. 2B). Differentially abundant taxa significantly correlated with the genotype included members of many leaf-colonizing phyla. Specifically, one-third of the differentially abundant taxa were Betaproteobacteria (order Burkholderiales, families Alcaligenaceae, Burkholderiaceae, Comamonadaceae, and Methylophilaceae), one-fourth were Alphaproteobacteria (Caulobacteraceae, Rhizobiaceae, and Rhodobacteraceae), and additional taxa belonged to Actinomycetota (Microbacteriaceae, Pseudonocardiaceae), Gammaproteobacteria (Moraxellaceae, Xanthomonadaceae), and Bacteroides (Spirosomaceae). Thus, the higher bacterial load in CLLF is not due to the enrichment of a single or a few dominant taxa, which would typically indicate dysbiosis, but instead reflects an apparently balanced enrichment of multiple bacterial families. Supporting this balance, CLLF plants never developed a disease phenotype in laboratory conditions (Fig. 1A). To further test whether CLLF’s microbiome composition contributes to plant fitness, we assessed its survival during overwintering in an outdoor garden experiment. CLLF showed a survival rate similar to NG2 (Fig. 2C). Additionally, leaf physiological functions in CLLF were only slightly altered compared to NG2. Specifically, the electron transport rate (ETR) and vapor pressure differential were slightly higher, while the stomatal conductance was slightly lower in CLLF (p < 0.01) (Supplementary Fig. 3). Together, these results suggest that CLLF recruits an alternative, but balanced, bacteriome in the same soil as NG2.

**Fig. 2.**
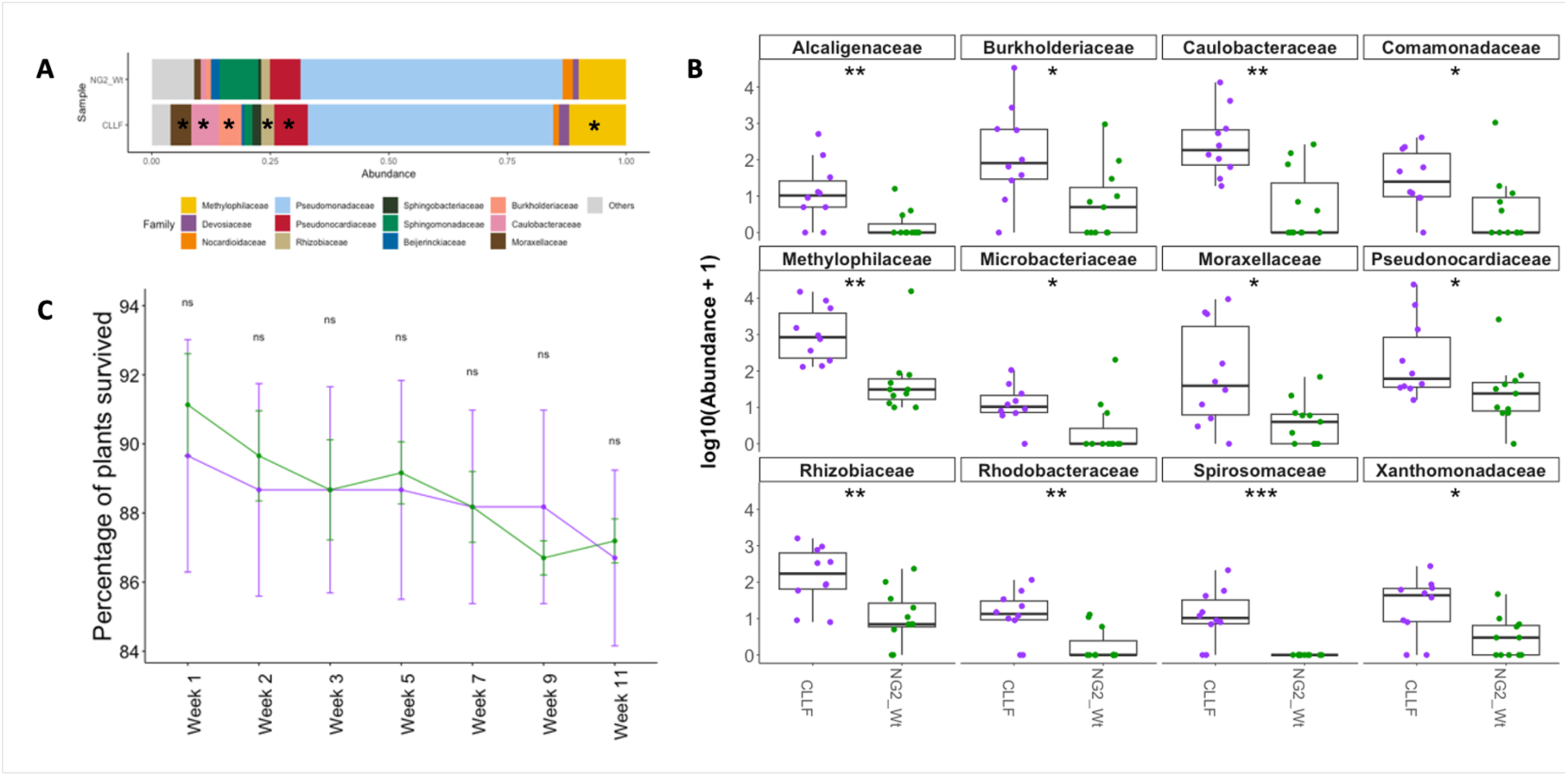
CLLF’s microbiome is not dysbiotic and it has normal survival rates. A) Relative abundance of the top 12 Families (remaining families represented as “Others”) in CLLF and NG2_Wt. Asterisks denote significant differences in abundance compared to the NG2 Wt. B) Differentially abundant bacterial families associated with genotypes. All families shown have a p-adjusted value <0.01 and data was normalized to plant-specific GI gene. C) Percentage of plants that survived from November 2022 to March 2023 in the garden (n=7). (CLLF; purple, NG2_Wt; green, ns; not significant, * p<0.05, ** p<0.01, *** p<0.001).

### CLLF’s immune system is intact and constitutively active

To test whether the CLLF immune system is intact, we investigated defense hormone levels. Notably, CLLF exhibited constitutively higher levels of defense hormones, with upregulated salicylic acid (SA) (Fig. 3B) and jasmonates in comparison to NG2 (Fig. 3A). CLLF plants grown in axenic conditions also showed higher levels of SA and jasmonates (Supplementary Fig. 4), indicating that the constitutive production of these hormones is not due to microbial colonization. Next, we analyzed CLLF gene expression relative to NG2. In agreement with the overproduction of defense hormones, gene expression profiles underscored CLLF’s constitutively active immune system. Genes involved in SA responses, systemic acquired resistance (SAR), and responses to bacterial molecules and oomycetes were among the most highly upregulated in CLLF (Fig. 3D). The strongest downregulated genes were primarily involved in glucosinolate-related processes. An overactive immune system could potentially be dysfunctional, which might explain higher commensal bacterial loads in CLLF. However, we found that CLLF remains capable of mounting a functional immune response. CLLF plants were more resistant to the bacterial pathogen *Pseudomonas syringae* pv. *syringae* DC3000 (Supplementary Fig. 5C) and demonstrated higher resilience to the fungal pathogen *Sclerotinia sclerotiorum* (Supplementary Fig. 5A and 5B) compared to NG2. These results suggest that plant defenses that maintain immunity to pathogens can be fully functional in parallel with the recruitment of non-pathogenic leaf bacterial communities.

**Fig. 3.**
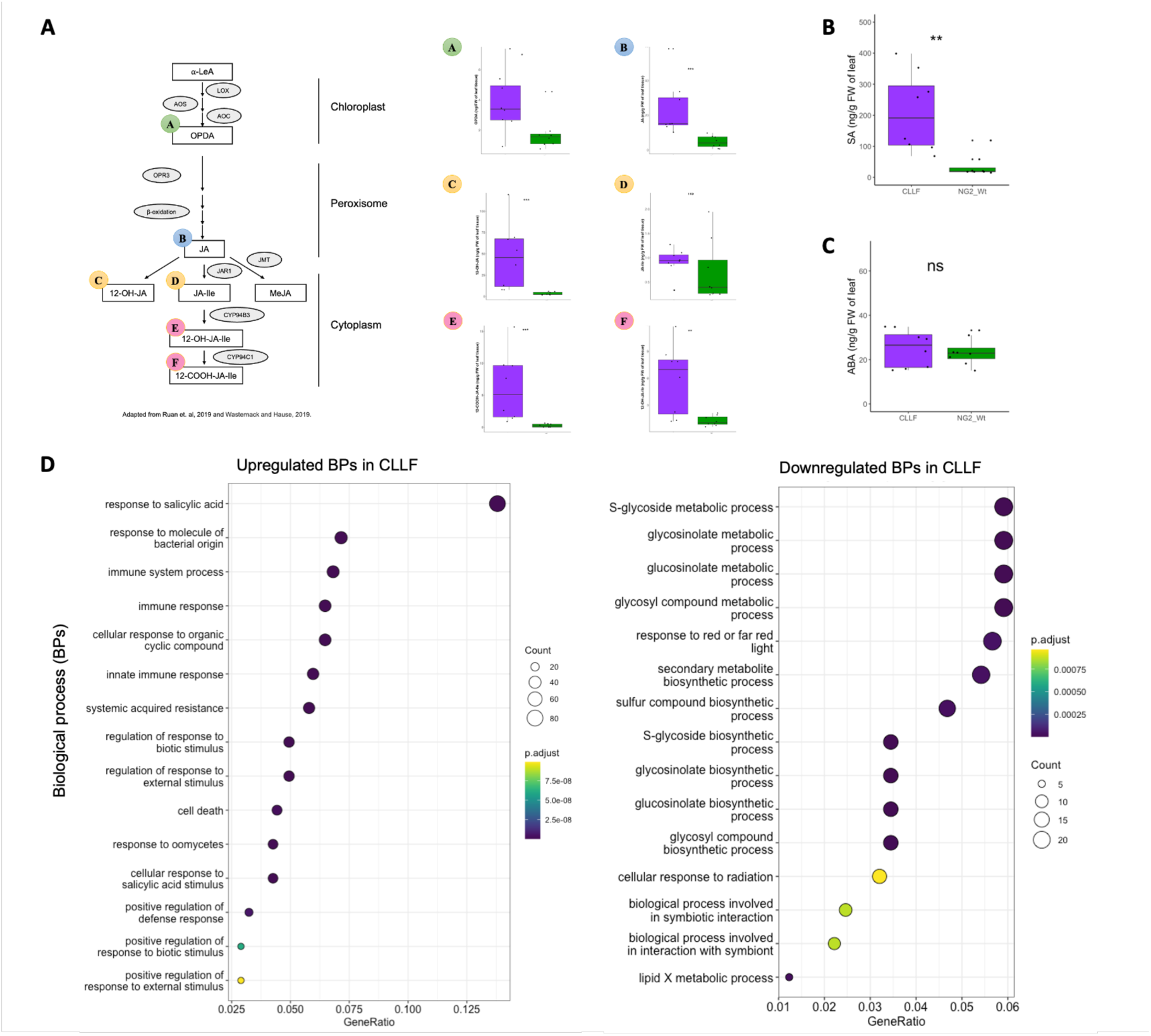
CLLF shows upregulated defense hormone profiles and has a constitutively active immune system. A) Jasmonic acid (JA) biosynthesis precursors and derivatives measured in nanograms per fresh weight of leaf tissue (n=8). A schematic representation of the JA biosynthesis pathway is shown next to it (Highlighted in color are each precursor/derivative). B) Total salicylic acid (SA) levels in CLLF and NG2 measured in nanograms per fresh weight of leaf tissue (n=8). C) Total abscisic acid (ABA) levels in CLLF and NG2 measured in nanograms per fresh weight of leaf tissue (n=8) (CLLF; purple, NG2; green, ns; not significant, * p<0.05, ** p<0.01, *** p<0.001). D) Gene ontology (GO) enrichment analysis of differentially expressed genes in gnotobiotic CLLF compared to the NG2. The cut-off for the genes was set at log fold change > [2], base mean > 100, and p-value < 0.01.

### Some leaf bacteria may directly utilize hormones as a resource for growth

High salicylic acid (SA) levels were previously linked to increased bacterial diversity in the phyllosphere (17), and SA has been shown to recruit specific taxa directly in the rhizosphere (9). Thus, we hypothesized that plant hormones may simultaneously shape plant immunity and recruit leaf bacteriomes, explaining the seemingly counterintuitive resistance and recruitment phenotypes of CLLF. To test this, we examined whether strains representative of diverse CLLF-associated phyla could utilize salicylic acid as a carbon source (Fig. 4). We found that some bacteria belonging to Alphaproteobacteria (Rhizobium A2 and Methylorubrum), Actinomycetes (Nocardia, Nocardioides, Brevibacterium, Rhodococcus) and Bacillota (*Bacillus A3*) showed growth in minimal media supplemented with SA as the sole carbon source. However, the Betaproteobacteria, Gammaproteobacteria, and Bacteroides strains we tested did not show growth in SA *in vitro*. This result is consistent with the idea that the overproduction of SA - and potentially other hormones - contributes to the recruitment of specific bacterial taxa that can metabolize them, supporting previous findings on the role of plant hormones in microbiome assembly.

**Fig. 4.**
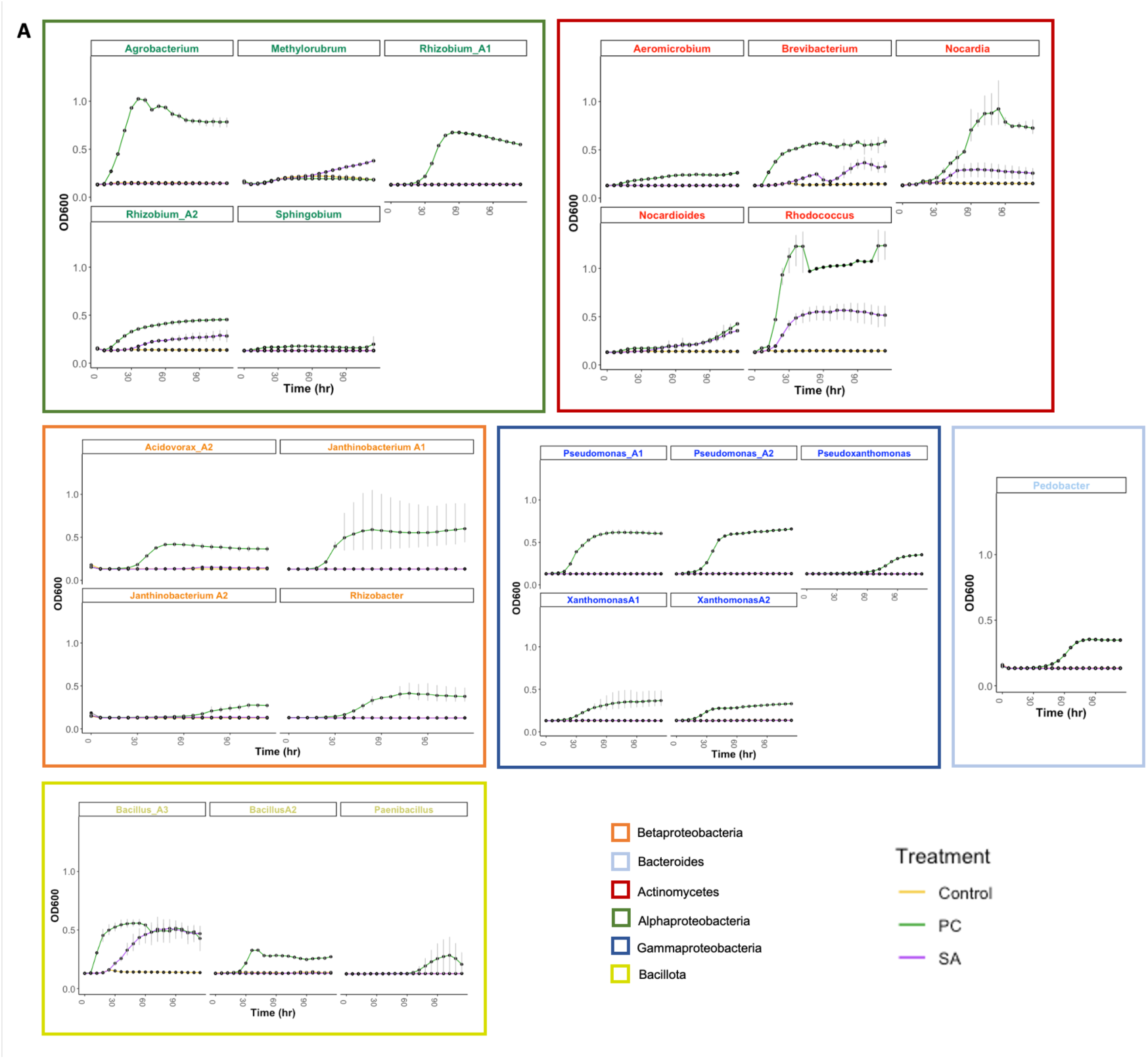
Growth rate of representative strains from various phyla in minimal media supplemented with salicylic acid (SA) as carbon source. Control shows growth in minimal media without any carbon source, colored yellow. PC (positive control) is growth in minimal media supplemented with sucrose and fructose (11 mM sucrose + 11 mM fructose), colored green. SA is the growth in minimal media supplemented with salicylic acid (0.5 mM Salicylic acid) as a carbon source, colored purple. The growth rate is measured as optical density at 600nm (n=4).

### High defense-related gene expression in CLLF does not decrease at elevated temperatures

A well-known effect of moderately high temperatures in plants is the downregulation of defense-related genes, leading to increased pathogen susceptibility. During an accidental breakdown of plant growth chambers, where temperatures reached 34°C, we observed that CLLF exhibited higher tolerance to extreme heat than NG2 (Supplementary Fig. 5D). This enhanced tolerance could be linked to increased SA signaling (18). To better understand this, we analyzed gene expression differences between CLLF and NG2 under both normal (18°C night, 22°C day) and high-temperature (30°C day and night) conditions (Supplementary Fig. 6). Overall, the gene expression profile of CLLF remained stable despite heat treatment (Supplementary Fig. 6A). Specifically, very few genes and associated biological processes were differentially regulated solely in response to heat stress (Supplementary Fig. 6B). In contrast, nearly all processes that were differentially regulated in CLLF under normal conditions compared to NG2 were similarly regulated after heat exposure (Supplementary Fig. 6C). This included immune activation-related processes, which are typically downregulated at elevated temperatures (11). Thus, the constitutively active immune system of CLLF maintains its activity regardless of temperature, potentially contributing to its increased heat tolerance.

### CLLF bacteriome imbalance causes dysbiosis at high temperatures, but does not affect NG2

Since CLLF’s activated immune system may be linked to its alternative bacteriome balance and did not exhibit the typical downregulation of immune responses at high temperatures, we investigated whether bacteriome composition influences balance during temperature stress. To address this, we conducted a semi-gnotobiotic synthetic community (SynCom) experiment (see methods), where plants were pre-inoculated with either a full SynCom representative of the entire CLLF bacteriome (SynCom1) or SynComs with major taxonomic groups removed. This approach allowed us to determine whether reducing taxonomic diversity affects plant stability. SynCom2 lacked Burkholderiales (Comamonadaceae and Oxalobacteriaceae), SynCom3 lacked Bacteroidota (Flavobacteriaceae and Sphingobacteriaceae), and SynCom4 lacked both groups (Supplementary Fig. 7 and Supplementary Table 1). Notably, bacteria remaining in SynCom4 included Actinomycetes, Bacillota, and Alphaproteobacteria, some of whom utilized salicylic acid as a carbon source (Fig. 4).

Plants not exposed to heat displayed normal, healthy phenotypes across all SynCom treatments (Fig. 5A and Supplementary Fig. 8A). To further investigate effects of microbiome composition, we evaluated gene expression profiles of NG2 and CLLF treated with SynCom1 and SynCom4 under normal temperatures. As expected, CLLF with SynCom1 exhibited upregulated immune responses compared to NG2, mirroring its response in both axenic and natural soil (Fig. 3D, Supplementary Fig. 6, Fig. 5B). In response to the reduced SynCom4, NG2 exhibited significantly increased immune responses, activation of phytoalexin biosynthetic pathways, and downregulation of photosynthesis-related genes, while CLLF maintained high defense-related gene expression (Fig. 5B and 5C). This suggests that removing Burkholderiales and Bacteroidota in SynCom4 required a defense response to maintain balance in NG2, while CLLF likely retained balance due to its already upregulated immune system.

**Fig. 5.**
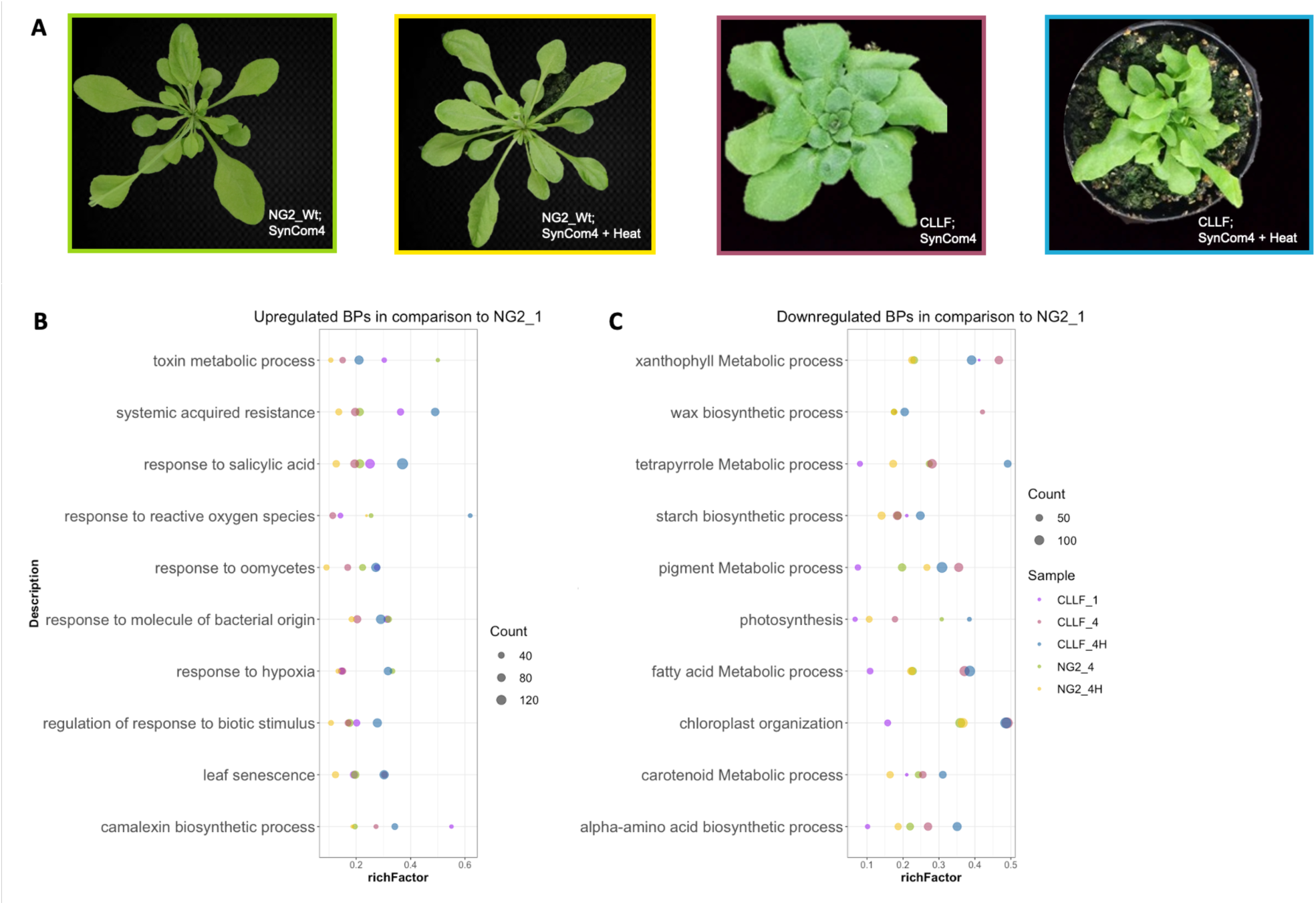
Phenotypic and gene expression responses of SynCom-inoculated plants under normal and heat stress conditions. CLLF exhibits dysbiosis at higher temperatures when inoculated with a taxonomically reduced community. (A) Phenotypic comparison of SynCom4-inoculated plants under normal conditions (18/22°C) and after heat stress (30°C for 48h). Light green border: NG2 inoculated with SynCom4 at normal temperature. Yellow border: NG2 inoculated with SynCom4 after heat stress. Pink border: CLLF inoculated with SynCom4 at normal temperature. Blue border: CLLF inoculated with SynCom4 after heat stress. (B and C) Upregulated (B) and downregulated (C) biological processes (BPs) across conditions. Comparison includes CLLF plants inoculated with SynCom1 (diverse community) at normal temperature (purple), NG2 and CLLF plants inoculated with SynCom4 at normal temperature (light green and pink, respectively), and NG2 and CLLF plants inoculated with SynCom4 after heat stress (yellow and blue, respectively), all relative to NG2 inoculated with SynCom1 at normal temperature. All genes used for biological process assignments met the criteria: p ≤ 0.01, base mean > 100, and fold change ≥ |2|. The rich factor represents the proportion of differentially expressed genes annotated to each term relative to background genes. The top 10 commonly upregulated and downregulated biological processes are shown, with additional BPs provided in Supplementary Fig. 9.

Upon exposure to 30°C for 48 hours, NG2 maintained a healthy phenotype regardless of SynCom treatment (Fig. 5 and Supplementary Fig. 8B). The immune responses previously activated by SynCom4 in NG2 under normal conditions were downregulated following heat stress, indicating that NG2 appropriately adjusted immune signaling to maintain stability, even with a highly reduced bacterial community. In contrast, CLLF with SynCom4 exhibited severe symptoms of heat stress, including deformed rosettes and chlorosis (Fig. 5A and Supplementary Fig. 8C). These plants displayed extremely high expression of immune-related genes, senescence markers, and low-oxygen response genes, suggesting a potential growth-defense tradeoff under stress. However, CLLF with SynCom1 showed no signs of dysbiosis after heat treatment (Supplementary Figs. 8 and 9), reinforcing the idea that a complete bacteriome helps maintain stability under stress conditions.

Together, these results suggest that when the bacteriome is incomplete, proper immune regulation is necessary to avoid dysbiosis under heat stress. In the absence of proper regulation, a complete bacteriome counteracts dysbiosis, highlighting the critical role of microbiome diversity in supporting host stability during abiotic stress. Notably, CLLF consistently exhibited higher resistance than NG2 to *Pseudomonas syringae* pv. *syringae* DC3000 (Pst DC3000), even when inoculated with SynCom4 and exposed to heat treatment (Supplementary Fig. 10). This further supports the conclusion that the observed growth-defense imbalance under heat stress is linked to microbiome assembly and that this functions independently of pathogen resistance.

## Discussion

The absence of key components of the immune system has been previously shown to lead to dysbiosis, resulting in a breakdown of a healthy plant microbiome (5, 7). In these cases, disease phenotypes were characterized by the overgrowth of specific taxa, suggesting that plant immune systems maintain balance by preventing the proliferation of opportunistic pathogens (2). Here, we identified CLLF, a genotype that recruits unusually high leaf bacterial loads when grown in natural soil. Detailed phenotyping revealed that CLLF has a constitutively active and apparently fully functional immune system despite high bacterial loads. Compared to previously reported dysbiotic bacteriomes in immune-deficient genotypes (5, 7), CLLF was healthy and robust, with high diversity in leaf-associated bacterial taxa and no overgrowth of specific opportunistic pathogens. These findings suggest that CLLF’s constitutively active immune system and associated high bacterial loads represent an alternative, balanced state of the leaf bacteriome. Although high bacterial loads are often associated with disease when opportunistic pathogens proliferate (19), balanced recruitment may allow plants to sustain higher microbial loads without negative consequences. Additionally, our results indicate that although intact immunity keeps commensal bacteria from becoming pathogenic, the recruitment of commensal bacteria to leaves is compatible with and may even be directly linked to an active immune system.

The observation that constitutively activated immunity correlated with increased bacterial recruitment in CLLF could be partially explained by the ability of certain bacteria to use plant defense signals as resources for growth. Some bacteria are already known to metabolize salicylic acid (SA) directly (20), and SA has been suggested to play a role in rhizosphere recruitment (9) as well as promoting leaf bacterial diversity (17). SA is also likely available to leaf colonizers, as it accumulates in extracellular compartments (e.g., the apoplast) upon immune activation (21). In our tests, the ability of bacterial taxa used in the SynCom experiments to grow on SA followed taxonomic patterns: none of the Betaproteobacteria or Bacteroidota tested showed growth in SA, while the tested Alphaproteobacteria, Actinomycetes, and Bacillota did. This suggests that SA may directly contribute to the recruitment of these taxa in the phyllosphere.

Apart from SA, CLLF also accumulated jasmonates, including OPDA and JA, at higher levels compared to NG2. Although this did not include JA-isoleucine, the best-studied bioactive form, molecules such as OPDA play key roles in defense, thermotolerance, and hormone regulation (22, 23). Additionally, JA species have been shown to affect plant-associated bacterial activity, acting as chemoattractants and biofilm inducers in the rhizosphere (24, 25). Furthermore, the aliphatic glucosinolate profile of CLLF was significantly altered compared to NG2 (data not shown). Since some bacteria metabolize glucosinolates, and glucosinolates are known to influence bacterial recruitment (10), these changes likely contributed to the distinct leaf microbiome composition in CLLF. Together, these findings suggest that secondary metabolites linked to plant immunity likely play a direct role in recruiting non-pathogenic leaf bacteria.

Alternative roles for plant hormones are not surprising, as they are ancient signals in the green lineage and likely serve multiple functions. For example, SA-responsive NPR proteins function as SA receptors in all plants, but their conserved roles in light and temperature sensing predate their function in defense, which likely evolved later (26). If plant defense signals also play a role in microbial recruitment, forward genetics approaches such as screening for genotypes with altered metabolite profiles could help engineer plant microbiomes in a targeted way. However, leveraging this strategy requires further research to determine how specific metabolites influence bacterial recruitment across different environmental conditions.

The SA pathway is integral to plant immunity, playing a major role in resistance to both biotic and abiotic stressors (26, 27). However, the downregulation of SA biosynthesis at high temperatures makes plants more vulnerable to pathogens (28). To counteract this, exogenous SA application and genetic approaches to maintain SA accumulation under heat stress have been suggested (18, 28). Despite these potential benefits, the broader consequences of altering SA regulation in relation to plant-microbe interactions remain unclear. In this study, CLLF exhibited constitutive SA-dependent immune upregulation, which remained high under heat stress. As expected, CLLF displayed higher resistance to pathogens and increased heat tolerance compared to NG2. However, when grown in a microbiome with reduced taxonomic diversity, CLLF experienced severe dysbiosis at elevated temperatures. One possible explanation is that high SA levels under heat stress promoted the overgrowth of SA-metabolizing bacteria, which were no longer balanced by other microbial colonizers. Consistent with this, CLLF plants pre-inoculated with a diverse bacteriome did not experience dysbiosis, indicating that microbial diversity contributes to resilience by stabilizing immune signaling under environmental stress.

In conclusion, our findings suggest that alternative balanced states exist within the phyllosphere microbiome. Specifically, plant immune activation alters microbial recruitment patterns without necessarily causing dysbiosis. This may be explained by plant hormones acting as both defense signals and microbial recruitment cues, influencing colonization by non-pathogenic bacteria. However, our findings also indicate that hormonal balance plays a key role in maintaining microbiome stability under stress conditions. Since natural microbiome assembly depends on environmental factors and chance colonization events (29), not all beneficial taxa may always be present in every plant. Given this risk, plants may have evolved to downregulate immune responses under heat stress in part as a strategy to prevent unintended microbial imbalances. As plant engineering strategies increasingly focus on improving stress tolerance to combat climate change (30), it is essential to consider how immune signaling alterations influence plant-microbe interactions. Ensuring microbial diversity during colonization may help mitigate unintended consequences and enhance plant resilience in dynamic environments.

## Conclusions

Our study reveals that immunity and microbial recruitment are parallel processes that together shape plant resilience to temperature stress. By characterizing the *Arabidopsis thaliana* genotypes CLLF and NG2, we demonstrate that constitutive immune activation can coexist with high bacterial loads and diversity without leading to dysbiosis. CLLF’s leaf bacteriome includes multiple taxa capable of metabolizing immune-related compounds such as salicylic acid, suggesting that defense hormones may serve dual roles in immunity and microbiome recruitment. Crucially, we show that the maintenance of a taxonomically diverse leaf microbiome is essential for plant stability under heat stress. While CLLF’s immune system remains active at elevated temperatures, the loss of microbial diversity, specifically, the exclusion of key phyllosphere colonizing bacterial taxa compromises resilience and results in dysbiosis. In contrast, NG2, which downregulates defense under heat, avoids dysbiosis regardless of microbiome composition. These findings suggest that balancing immune signaling with microbiome diversity is key to plant health under abiotic stress. Our results underscore the importance of microbiome composition in shaping plant responses to environmental changes and highlight new avenues for enhancing crop resilience through targeted manipulation of plant-microbe interactions.

## Materials and Methods

### Plant genotypes

The *Arabidopsis thaliana* ecotype NG2 (NASC ID N2110865; Je-1) is a previously described genotype that assembles a characteristic bacteriome when grown in its native soil in Jena, Germany (10). For this study, we aimed to isolate a genotype that assembles an alternative balance in the natural soil compared to NG2. To do so, we chemically mutagenized NG2 seeds with ethyl methanesulfonate and grew them in the natural soil. Most of the mutant pool looked and behaved like NG2, but we isolated CLLF which clearly shows an unusual but healthy mutant phenotype and high bacterial loads. Genome sequencing and phylogenetic analysis of SNPs, however, suggest that CLLF diverges signficantly from NG2. CLLF is also significantly diverged from other local genotypes that we work with. In addition, NG2 genomes sequenced from three seed stock batches collected years apart are uniform, so seed stock contamination is unlikely (Supplementary Fig.1). Thus, we hold the most likely scenario to be that CLLF derives from an unintended cross of an NG2 mutant with a wild seed growing in the natural soil. This could be consistent with constitutively upregulated immunity, which can occur when incompatible resistance gene alleles are crossed. Although the exact origins of the CLLF genotype are therefore unknown, we decided to take advantage of its unique microbiome assembly phenotype for comparison to NG2. CLLF is named for its distinct phenotype, including <u>c</u>urled and compressed leaves, late flowering, and increased bacterial recruitment, making it an interesting genotype for studying host-microbiome interactions. For this study, all experiments were performed using seeds from the F4 generation of CLLF.

### Variant Analysis of Plant Genomes

*A. thaliana* genotypes isolated from nearby Jena, DE (NG2 (from two separate batches), JT1, PB, SW1 and Woe) were grown in 2020. NG2 as well as CLLF and the reference genotype Col-0 were grown in 2023. Leaf material was harvested from all genotypes and genomic DNA was extracted and sequenced on an Illumina HiSeq instrument (MiGs) in 2020 and on Illumina NovaSeq 6000 (SeqCenter) in 2023. After quality filtering to remove adapters, reads were mapped to the *Arabidopsis thaliana* Col-0 reference genome (TAIR10) (31) using bwa. Variants were jointly called using the recommended GATK4 pipeline for short variant discovery (32) implemented using snakemake wrappers version 1.12.2 (33). In short, variants were called per-sample using HaplotypeCaller, consolidated using GenomcsDBImport, and genotyped together using GenotypeGCVFs. Then the variants were recalibrated using VariantRecalibrator using the Arabidopsis 1001 genomes variants (34) as a training dataset and filtered using ApplyVQSR with the truth sensitivity filter level set to 99.5. Only SNPs were used to make a maximum likelihood phylogenetic tree using the package SNPhylo (35).

### Plant growth conditions

For microbiome analysis of plants colonized from the soil, seeds of the *A. thaliana* NG2 and the genotype CLLF were surface sterilized by washing for 2 min in 2% bleach followed by 30s in 70% ethanol and two sterile water rinses. They were sown onto commercial soil supplemented with garden soil inoculum after vernalization in 0.1% agarose at 4°C for 5 days. Later, a plastic wrap was used to cover individual pots, leaving a small hole in the middle to allow the seedling to germinate. This acted as a barrier between the leaves and soil, preventing detection of any soil microbes on leaves due to increased soil contact in CLLF (Barrier Experiment). Plants were then allowed to grow at 22°C day (16h) / 18°C night (8h) for 3 weeks in a plant climate chamber (PolyKlima, Freising, Germany) before leaf harvest. Each sample represents 4 leaves from a single plant. No plastic barrier was used for replicate experiments in which leaves were used either for total microbiome or phytohormone analysis. For phytohormone analysis, commercial soil without garden soil inoculum was used. For gnotobiotic growth, surface sterilized and vernalized seeds were sowed into Linsmaier and Skoog media (4.3g/L LS salt mix (36), 1% sucrose, 0.5% MES, 0.3% phytagel) solidified in sterile 24 well plates.

### Leaf bacterial load estimation

To compare bacterial loads between genotypes, leaf bacteriome composition was assessed using CFU counting. Seeds of NG2 and CLLF were surface sterilized and sown into individual pots containing commercial soil supplemented with a natural soil inoculum (60 g of garden soil dissolved in 1L autoclaved Milli-Q water, mixed with 3L commercial soil). Plants were grown under controlled conditions (22°C day (16h) / 18°C night (8h) cycle) on growth shelves. Once the plants reached 3.5 weeks of age, two leaves per plant were collected for **c**olony-forming unit (CFU) counting. Leaves were ground in sterile 1x PBS using bead beating (1400 rpm for 30s), and dilution series were prepared and plated onto R2 agar. Plates were incubated for 48 hours before CFU counting.

### Plant DNA extraction

We used a high throughput DNA extraction method to isolate DNA from leaves. 2-6 leaves were taken for DNA extraction depending on the weight and after washing away dust and debris with autoclaved Milli-Q water, were frozen at -80°C. Later, frozen samples were ground in a bead beater at 1400rpm, for 30s to 60s to make a fine homogenate using 3mm metal and 0.25-0.5 mm glass beads. DNA was extracted in CTAB (2% (w/v) Cetylimethylammoniumbromid, 100mM Tris with pH 8.0, 20mM EDTA with pH 8, 1,4M NaCl, 1% (w/v) Polyvinylpyroledon) buffer and then purified from the lysate using Phenol:Chloroform:Isoamyl alcohol (25:24:1) and precipitated using 0.7 volume 2-propanol. Precipitated DNA was then dissolved in 10mM-Tris buffer for the barrier experiment or nuclease-free water for the remaining microbiome analysis.

### 16S rRNA gene amplicon sequencing

For amplicon library preparation, we used a modified version of the host-associated microbe PCR (hamPCR) protocol (16). Kapa HiFi enzyme (Roche) was used for all PCR reactions. Zymomix, nuclease-free water, and CTAB extraction buffer were used as internal controls to evaluate contamination and sequencing depth. In the first tagging step, we used universal primers targeting the V3-V4 region of 16S rDNA modified with a 5’ overhang. In addition to that, we used primers targeting host-specific single copy gene (GIGANTEA or GI gene) and blocking oligos to limit the amplification of chloroplast 16S rDNA (37). Each 10 µL 1^st^ PCR mix contained 1x Kapa Buffer, 0.3 mM Kapa dNTPs, 0.08 μM of each of the forward and reverse 16S rDNA and GI primers, 0.25 μM of each of the blocking oligos, 0.2μL Kapa HiFi DNA polymerase, and 50 to 100 ng template genomic DNA. Thermocycling steps included initial denaturation at 95°C for 3min, denaturation at 98°C for 20s, two-step annealing at 58°C for 30s and 55°C for 1min, extension at 72°C for 1min (cycle repeated 5x), and final extension at 72°C for 2min. An enzymatic cleanup was done to remove primer dimers and inactivate nucleotides by directly adding 0.5µL of Antarctic phosphatase and Exonuclease 1 (New England Biolabs, Inc) and 1.22µL Antarctic phosphatase buffer (1x final concentration) to the 1^st^ PCR mix. It was then incubated at 37°C for 30 minutes followed by 80°C for 15 minutes to inactivate the enzyme activity. Next, in a second PCR barcoded primers targeted the 5’ overhangs to amplify 1^st^ PCR products. The 20µL 2^nd^ PCR mix contained 1x Kapa Buffer, 0.375 mM Kapa dNTPs, 0.3 μM of each forward and reverse primer, 0.5μL Kapa HiFi DNA polymerase, and 5µL 1^st^ PCR product. The PCR program included initial denaturation 95°C for 3min, denaturation at 98°C for 20s, annealing at 60°C for 1min, extension at 72°C for 1min (repeated 35x), and final extension at 72°C for 2min. The barcoded PCR products from the second PCR were cleaned by magnetic separation using Sera mag beads. Later, we estimated the concentration of each sample by measuring fluorescence using PicoGreen (Quant-iT^TM^ PicoGreen^TM^). After adjusting the concentration, all samples were pooled into one single library and concentrated using sera mag beads. The concentrated library was then loaded into a 2% agarose gel and stained with Roti gel stain. The separated GI and 16S rDNA bands were cut out of the gel and DNA was eluted from each of the gel pieces using the GeneJET gel extraction kit (Thermo Scientific). After measuring the concentration of gel-eluted DNA, a final library was prepared with an adjusted concentration of 95% 16S rDNA and 5% GI. We quantified the final library with a Qubit (Thermo Fisher Scientific, Inc). Then the library was denatured and loaded onto a MiSeq lane spiked with 10% PhiX genomic DNA to ensure sequence diversity. Using conventional Illumina sequencing primers, 600 cycles of Illumina sequencing were carried out to obtain 300 bp sequences in both the forward and reverse directions.

### Amplicon sequencing data processing

For all datasets, adapter sequences were first removed from reads using Cutadapt 3.5 and reads were split into samples according to barcodes using a custom script. Quality filtering and clustering the data into amplicon sequencing variants (ASVs) were performed with the dada2 (version 1.18.0) algorithm, using only forward reads because of its higher quality. Taxonomy was assigned to the ASVs using dada2 with the Silva 16S rDNA database (version 138.1) supplemented with the A. thaliana GI gene sequence. We examined positive and negative controls from all data sets. Positive control (Zymomix) had an expected distribution of taxa and negative controls had very few reads (<60 reads) suggesting minimal contaminations, so these were not processed further. R packages Phyloseq (version 1.34.0) (38), VEGAN (version 2.5-7)(39), and DEseq2 (version 1.44.0)(40), were used for downstream analysis. Host-derived reads were removed from the ASV tables by removing family “Mitochondria”, order “Chloroplast” and Genus “Arabidopsis GI”. In addition, all samples were normalized to GI reads before analysis so that diversity patterns reflect the true abundances of leaf bacteria. Plant GI reads were also used to normalize the total number of bacterial reads to get an estimate of the relative bacterial loads of each sample. Other packages such as ggplot2, reshape2, and ggpubr were used for statistical analysis and plotting data. Scripts for generating the main figures from the ASV tables and metadata will be made publicly available on Figshare prior to publication.

### Assessing plant survival rate in a garden experiment

To compare survival rates of CLLF and NG2, we germinated seeds in a mix of 1:4 garden soil to commercial soil (as described above) under laboratory conditions. Plants were sown into 7 replicate trays, each divided into two halves, one for each genotype. Plants were thinned to 30 seedlings of each genotype per tray, which was considered as one replicate. The trays with seedlings were moved to the garden 7 days after germination (in Nov 2022), and the number of plants surviving in each tray was counted every week until plants started flowering (in March 2023).

### Phytohormone analysis

For the phytohormone extraction, we used a previously published high-throughput method (41). We collected 100mg of leaves from 3-week-old lab-grown plants in a pre-weighed 2mL screw-cap tube with two metal beads each. Then, we fast-froze the samples in liquid nitrogen and stored at -80°C until further processing. The frozen samples were ground into a fine powder using a tissue lyser (MiniG 1600, SPEX Sample Prep). For the extraction, samples were homogenized in 15:4:1 (v/v/v) methanol: H2O: formic acid extraction buffer containing isotope-labeled phytohormone standards, centrifuged at high speed, and the supernatant was collected. The supernatant was then passed through a reversed-phase solid-phase extraction (SPE) column (HR-X, Machery-Nagel, 738530.025M). Methanol from the flow-through was then allowed to evaporate and the remaining samples were reconstituted in 1N formic acid. Reconstituted samples were then passed through a mixed-mode cation exchange SPE column (HR-XC, Machery-Nagel, 738540.025M), and the flow through was discarded. The column was washed once with 1N formic acid and eluted using an 80% aqueous methanol containing 0.2N formic acid. This eluted fraction contained abscisic acid (ABA), salicylic acid (SA), and jasmonates (JAs), which were analyzed by LC-MS/MS on a triple-quadrupole mass spectrometer (EVO-Q Elite, Bruker) using the chromatog. The chromatographic condition and multiple-reaction-monitoring details for the compound detection are described in detail by Schäfer et al. (41).

### Pathogen infection assays

For the bacterial pathogen infection assay, CLLF and NG2 were grown in gnotobiotic conditions in LS media in square petri dishes. *Pseudomonas syringae pv syringae* DC3000 (Pst DC3000) was grown in LB agar supplemented with rifampicillin for 48 h at 30°C. Colonies were then collected using a loop and resuspended in 10 mM MgCl2. OD was measured using a UV spectrometer and adjusted to 0.02 OD. 0.02 % Silwet L-77 was freshly added to the bacterial suspension just before flood inoculation. For flood inoculation, 40mL bacterial culture was poured into the petri dishes with plants. Plants inoculated with 40mL of 10 mM MgCl2 plus 0.02 % Silwet L-77 were taken as controls. Disease symptoms were measured after 24h.

For the fungal infection assay, plants grown in the semi-gnotobiotic (2-week-old seedlings grown in LS media transplanted into commercial soil) system. *Sclerotinia sclerotiorum* (Ssc) was grown in Potato-Dextrose agar (PDA) plates from a subculture for 4-6 days. We collected agar discs from the actively growing region of Ssc plates and used this to inoculate liquid PDB (Potato-Dextrose Broth). Liquid culture was then allowed to grow overnight at room temperature with shaking. We harvested the mycelia the next day after passing it through an autoclaved nylon membrane. After washing them with autoclaved milliQ water, they were resuspended in fresh PDB. This was then macerated to get small hyphal fragments using a sterile macerator. Hyphal fragments were adjusted to 2x10^5^ fragments/ml. 5µL of this hyphae suspension was used to inoculate leaves of 4-week-old plants. The leaf midrib was avoided for easy measurement of the lesion and lesion size was measured at both 18 hours post-inoculation (hpi) and 40hpi.

### Synthetic community preparation

We used bacterial strains that were previously isolated from CLLF or NG2 leaves for preparing the Synthetic Communities (SynComs) (Supplementary Table 1A). All strains were grown on R2 agar plates. Depending on each strain’s growth rate, it took 1 to 5 days to grow all strains. The bacteria were then scraped off the plates and resuspended in 1x phosphate buffer saline (PBS). Optical density (OD) was measured using a 1:10 dilution of each bacterial suspension in a plate reader. Since we observed huge variation in the number of CFUs per OD of different bacteria, we decided to consider CFUs for preparing SynComs. Therefore, an OD to CFU relationship of each bacterium was measured and this was used to combine the bacterial suspensions into a single inoculum so that all strains would have approximately same number of CFUs (∼30 CFU per strain). A total of 45 strains belonging to diverse plant-associated bacterial phyla constituted the SynCom1. The SynCom2 lacked strains belonging to Comamonadaceae and Oxalobacteriaceae (Betaproteobacteria). SynCom3 lacked strains belonging to

Bacteroides and SynCom4 lacked Betaproteobacteria and Bacteroides. For each missing strain in SynComs 2-4, we added the same amount of 1x PBS buffer to the SynCom stocks. In addition, aliquots of all four SynComs were mixed with an equal amount of 40% glycerol for future analysis. To check the SynCom compositions, 12µL from the glycerol stocks were later inoculated onto R2 agar plates and grown for 5 days at 28°C. All cells were harvested from R2 plates by suspending into 1x PBS, and later, all samples were centrifuged at 5000rpm in a table-top centrifuge and the pellet was resuspended in CD1 solution of DNAeasy plant pro kit (Qiagen; 0142924730) along with 0.25-0.5 mm glass beads. It was then bead-beated at 1400rpm for 30s. DNA was extracted using the kit following the suggested protocol. The communities were analyzed using the 16S rRNA gene amplicon sequencing pipeline as described above.

### Semi-gnotobiotic plant growth conditions and stress treatments

We noted that in fully gnotobiotic, enclosed growth systems, plants tended to become stressed by humidity. Therefore, we developed a semi-gnotobiotic system in which plants are pre-colonized by a defined community in a gnotobiotic environment and then moved to a standard, non-sterile laboratory soil and grown in trays. When we tested the semi-gnotobiotic system with a fluorescently labeled isolate of *Stenotrophomonas* sp., leaf microbiomes were dominated by *Stenotrophomonas* even after several weeks in the soil, suggesting that this is an effective inoculation strategy (42). For this study, surface-sterilized and vernalized seeds of CLLF and NG2 were sown into LS media. 7-day-old seedlings were inoculated with 5µL inoculum of one of the four SynComs or 1x PBS buffer. This was enough inoculum to wet the leaves and roots. There were 48 replicates per genotype and SynCom. The plants were grown for 1 week in LS before being transplanted into commercial soil at 22°C day-16h/18°C night-8h cycles. After 2 weeks of normal growth in soil, 16 plants from each genotype inoculated with one of the 4 SynCom or 1x PBS were given heat stress by moving into a growth chamber set at 30°C day (16h) nd night (8h) for 48h. The remaining plants were allowed to grow at normal 22°C day-16h/18°C night-8h cycles. Later the plants that underwent heat stress were moved back into normal growth conditions and all plants were grown for 2 days under 22°C day-16h/18°C night condition before treating with *Pseudomonas syringae pv. syrinage* DC3000 (Pst DC3000). Half of the plants from each treatment were spray-inoculated with 0.02 OD cultures of Pst DC3000 suspended in sterile milliQ water supplemented with 0.02% Silwet L-77 and the remaining half with sterile milliQ water. After 48 hours post-inoculation, we measured leaf function using an LI-COR 600, and 2-3 leaves from at least 6 plants were sampled for microbiome analysis from all the different treatments. 2-3 leaves from SynCom 1, 4, and negative controls with and without heat stress were sampled for RNAseq analysis (Method Fig. 1).

**Method Fig.1.**
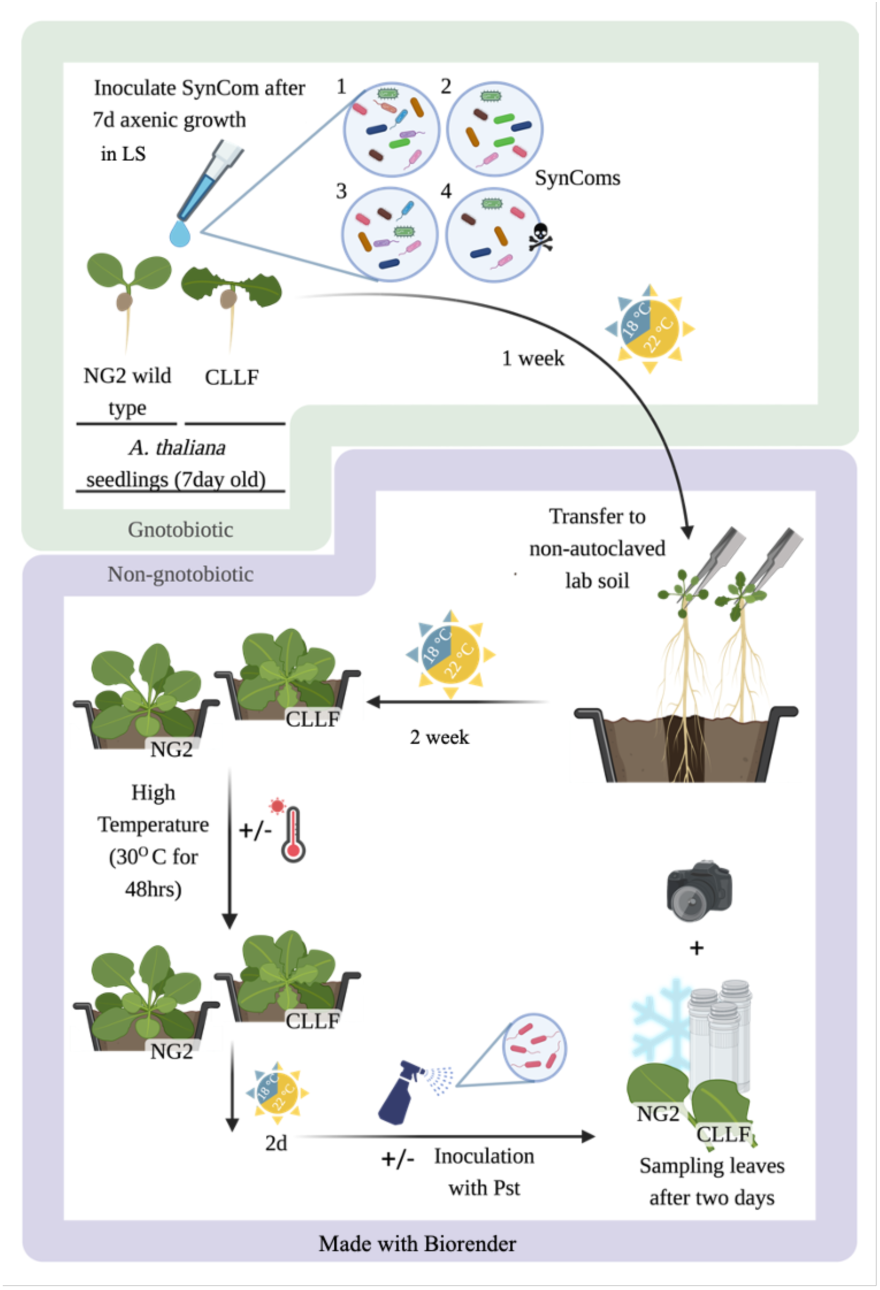
Workflow showing SynCom experiment in a semi-gnotobiotic system. Gnotobiotic phase; Sterilized seeds of CLLF and NG2 plants were germinated in sterile LS media. 7-day old seedlings were inoculated with 5µL SynCom or 1X PBS. Plants were allowed to grow in LS media with or without the SynComs for 1 week at 22h day/8h night cycles. **Non-gnotobiotic phase;** 2-week-old pre-colonized plants are moved to commercial soil. After two weeks of growth (22h day/8h night cycles), half the of plants were challenged with temperature stress at 30°C for 48 h, followed by 2 days of recovery under nomal condition. Later half of plants from each treatment was challenged with PstDC3000. After 2 days, leaves were sampled for microbiome or RNAseq analysis.

### RNA sequencing and data processing

All samples for RNAseq were fast-frozen using liquid nitrogen and stored at -80°C. The samples were then ground to a fine powder in a bead beater using 2 metal beads. Finely ground samples were used for RNA isolation using the RNAeasy Plant Mini Kit (Qiagen, ID: 74904). Total RNA was sent to Eurofins genomics for rRNA depletion, cDNA synthesis, library preparation and sequencing. Raw data received from Eurofins Genomics were then used for downstream processing as follows. Raw sequencing reads were assessed for quality using FastQC (version 0.11.9; https://www.bioinformatics.babraham.ac.uk/projects/fastqc). Adaptor trimming, quality filtering, and read preprocessing were performed using fastp (version 0.23.2) (43). In detail, 5′ and 3′ bases with a Phred quality score below 28 were cut and reads were removed if they had more than one ambiguous base, an average quality score below 28, or a length of fewer than 15 bases. Processed reads were aligned to the current *A. thaliana* genome (tair10.1) using Hisat2 (version 2.2.1) with standard parameters (44). The aligned reads were sorted and indexed using SAMtools (version 1.11) (45). Read counting was performed using featureCounts (version 2.0.1) (46) with the tair10.1 annotation as a reference. Differential gene expression analysis was performed using DESeq2 (version 1.38.3) (40) and comparisons having a false discovery rate (FDR) adjusted p-value <0.05 were deemed to be statistically significant. Gene ontology (GO) enrichment was performed for each DE gene set using the R package, clusterProfiler (version 4.10.0), and the GO category was assigned using the annotation data package org.At.tair.db (version 3.18.0). The detailed scripts with step-by-step instructions were uploaded to GitHub (https://github.com/Bioinformatics-Core-Facility-Jena/SE20231212_167).

### Growth assay with salicylic acid

All strains belonging to Betaproteobacteria and Bacteroides and representative strains from the remaining major taxonomic groups of the SynComs were used for growth assays. First, bacteria were grown in R2 agar at 28°C for 4 days. Fresh colonies were then harvested from agar plates using a sterile loop and resuspended in 1x PBS by vortexing and repeated pipetting. After measuring the optical density (OD_600_), the OD_600_ of all samples was adjusted to 0.02. 10 µL of this pre-adjusted culture was used as an inoculum for measuring growth in 190 µL of M9 minimal media (M9 salts: 33.7mM Na_2_HPO_4,_ 22mM KH_2_PO_4_, 8.55mM NaCl, 9.35mM NH_4_Cl, and 2 mM MgSO_4_*7H_2_0, 0.2 mM CaCl_2_, 1 μg/mL biotin, 1 μg/mL thiamine, and trace elements: 134 μM EDTA, 31 μM FeCl_3_-6H_2_O, 6.2 μM ZnCl_2_, 0.76 μM CuCl_2_-2H_2_O, 0.42 μM CoCl_2_-2H_2_O, 1.62 μM H_3_BO_3_, 0.081 μM MnCl_2_-4H_2_O) supplemented either with 0.5mM Salicylic acid (SA) as in (9), 11 mM Fructose plus 11 mM Sucrose (PC) or no additional carbon source (Control) for the growth assay. All samples were then incubated at 28°C with shaking (220rpm) in 15-minute intervals in a plate reader for 5 days. OD_600_ was measured every 1.5 h.

## Data Sharing Plan

Data needed to evaluate the conclusions in the paper are present in the paper and/or the Supplementary Materials. Raw sequencing data is available on NCBI-SRA for 16S data: PRJNA1120601 and RNAseq data: PRJNA1141903 will be released prior to publication). Processed data with code to generate the main figures will also be available on Figshare before final publication. The detailed scripts with step-by-step instructions for RNA-seq analysis were uploaded to GitHub (https://github.com/Bioinformatics-Core-Facility-Jena/SE20231212_167).

## Funding Information

Carl Zeiss Foundation via Jena School for Microbial Communication (JJ, MTA)

Deutsche Forschungsgemeinschaft (DFG, German Research Foundation) under Germany’s Excellence Strategy - EXC 2051 – Projekt-ID 390713860 (JJ, ET, TM, MTA, MM)

Max Planck Society (RH)

The Ministry for Economics, Sciences and Digital Society of Thuringia (TMWWDG), under the framework of the Landesprogramm ProDigital [DigLeben-5575/10-9]. (EB)

Deutsche Forschungsgemeinschaft (DFG, German Research Foundation) under CRC 1076 ‘AquaDiva’, subproject A06. (MM)

## References

1. McNew, George L., “The nature origin and evolution of parasitism” in Plant Pathology: An Advanced Treatise, (Academic Press, 1960), pp. 19–69.

2. F. Entila, X. Han, A. Mine, P. Schulze-Lefert, K. Tsuda, Commensal lifestyle regulated by a negative feedback loop between *Arabidopsis* ROS and the bacterial T2SS. Nat. Commun. 15, 456 (2024).

3. L. Maignien, E. A. DeForce, M. E. Chafee, A. M. Eren, S. L. Simmons, Ecological succession and stochastic variation in the assembly of *Arabidopsis thaliana* phyllosphere communities. mBio 5 (2014).

4. K. M. Meyer, et al., Plant neighborhood shapes diversity and reduces interspecific variation of the phyllosphere microbiome. ISME J. 16, 1376–1387 (2022).

5. T. Chen, et al., A plant genetic network for preventing dysbiosis in the phyllosphere. Nature 580, 653–657 (2020).

6. P. J. P. L. Teixeira, et al., Specific modulation of the root immune system by a community of commensal bacteria. Proc. Natl. Acad. Sci. 118, e2100678118 (2021).

7. S. Pfeilmeier, et al., The plant NADPH oxidase RBOHD is required for microbiota homeostasis in leaves. Nat. Microbiol. 6, 852–864 (2021).

8. C. J. Harbort, et al., Root-secreted coumarins and the microbiota interact to improve Iron nutrition in *Arabidopsis*. Cell Host Microbe 28, 825–837.e6 (2020).

9. S. L. Lebeis, et al., Salicylic acid modulates colonization of the root microbiome by specific bacterial taxa. Science 349, 860–864 (2015).

10. K. Unger, et al., Glucosinolate structural diversity shapes recruitment of a metabolic network of leaf-associated bacteria. Nat. Commun. 15, 8496 (2024).

11. B. Huot, et al., Dual impact of elevated temperature on plant defence and bacterial virulence in *Arabidopsis*. Nat. Commun. 8, 1808 (2017).

12. J. Qiu, et al., Warm temperature compromises JA-regulated basal resistance to enhance *Magnaporthe oryzae* infection in rice. Mol. Plant 15, 723–739 (2022).

13. H.-G. Mang, et al., Abscisic acid deficiency antagonizes high-temperature Inhibition of disease resistance through enhancing nuclear accumulation of resistance proteins SNC1 and RPS4 in *Arabidopsis*. Plant Cell 24, 1271–1284 (2012).

14. A. Issa, et al., Impacts of *Paraburkholderia phytofirmans* strain PsJN on tomato (*Lycopersicon esculentum* L.) under high temperature. Front. Plant Sci. 9, 1397 (2018).

15. R. J. Rodriguez, et al., Stress tolerance in plants via habitat-adapted symbiosis. ISME J. 2, 404–416 (2008).

16. D. S. Lundberg, et al., Host-associated microbe PCR (hamPCR) enables convenient measurement of both microbial load and community composition. eLife 10, e66186 (2021).

17. S. A. Vincent, A. Ebertz, P. D. Spanu, P. F. Devlin, Salicylic acid-mediated disturbance increases bacterial diversity in the phyllosphere but Is overcome by a dominant core community. Front. Microbiol. 13, 809940 (2022).

18. L.-J. Wang, et al., Salicylic acid alleviates decreases in photosynthesis under heat stress and accelerates recovery in grapevine leaves. BMC Plant Biol. 10, 34 (2010).

19. T. L. Karasov, et al., The relationship between microbial population size and disease in the Arabidopsis thaliana phyllosphere. [Preprint] (2019). Available at: http://biorxiv.org/lookup/doi/10.1101/828814 [Accessed 13 May 2024].

20. K. Ogata, M. Ohsugi, M. Tomita, T. Tochikura, The production of α-ketoglutaric acid from salicylic acid by *Pseudomonas* sp. Agric. Biol. Chem. 34, 364–369 (1970).

21. G.-H. Lim, et al., Plasmodesmata localizing proteins regulate transport and signaling during systemic acquired immunity in plants. Cell Host Microbe 19, 541–549 (2016).

22. W. Liu, S.-W. Park, 12-oxo-phytodienoic acid: A Fuse and/or switch of plant growth and defense responses? Front. Plant Sci. 12, 724079 (2021).

23. I. Monte, et al., An ancient COI1-independent function for reactive electrophilic oxylipins in thermotolerance. Curr. Biol. 30, 962–971.e3 (2020).

24. M. Antunez-Lamas, et al., Bacterial chemoattraction towards jasmonate plays a role in the entry of *Dickeya dadantii* through wounded tissues. Mol. Microbiol. 74, 662–671 (2009).

25. O. S. Kulkarni, et al., Volatile methyl jasmonate from roots triggers host-beneficial soil microbiome biofilms. Nat. Chem. Biol. 20, 473–483 (2024).

26. H. Jeon, et al., Contrasting and conserved roles of NPR pathways in diverged land plant lineages. New Phytol. 243, 2295–2310 (2024).

27. Y. Liu, et al., Diverse roles of the salicylic acid receptors NPR1 and NPR3/NPR4 in plant immunity. Plant Cell 32, 4002–4016 (2020).

28. J. H. Kim, et al., Increasing the resilience of plant immunity to a warming climate. Nature 607, 339–344 (2022).

29. M. R. Wagner, et al., Host genotype and age shape the leaf and root microbiomes of a wild perennial plant. Nat. Commun. 7, 12151 (2016).

30. B. N. Archibald, V. Zhong, J. A. N. Brophy, Policy makers, genetic engineers, and an engaged public can work together to create climate-resilient plants. PLOS Biol. 21, e3002208 (2023).

31. L. Reiser, et al., The *Arabidopsis* information resource in 2024. GENETICS 227, iyae027 (2024).

32. R. Poplin, et al., Scaling accurate genetic variant discovery to tens of thousands of samples. [Preprint] (2017). Available at: http://biorxiv.org/lookup/doi/10.1101/201178 [Accessed 28 March 2025].

33. F. Mölder, et al., Sustainable data analysis with Snakemake. F1000Research 10, 33 (2021).

34. C. Alonso-Blanco, et al., 1,135 genomes reveal the global pattern of polymorphism in *Arabidopsis thaliana*. Cell 166, 481–491 (2016).

35. T.-H. Lee, H. Guo, X. Wang, C. Kim, A. H. Paterson, SNPhylo: a pipeline to construct a phylogenetic tree from huge SNP data. BMC Genomics 15, 162 (2014).

36. E. M. Linsmaier, F. Skoog, Organic growth factor requirements of tobacco tissue cultures. Physiol. Plant. 18, 100–127 (1965).

37. T. Mayer, et al., Obtaining deeper insights into microbiome diversity using a simple method to block host and nontargets in amplicon sequencing. Mol. Ecol. Resour. 21, 1952–1965 (2021).

38. P. J. McMurdie, S. Holmes, phyloseq: an R package for reproducible interactive analysis and graphics of microbiome census data. PLoS One 8, e61217 (2013).

39. P. Dixon, VEGAN, a package of R functions for community ecology. J. Veg. Sci. 14, 927– 930 (2003).

40. M. I. Love, W. Huber, S. Anders, Moderated estimation of fold change and dispersion for RNA-seq data with DESeq2. Genome Biol. 15, 550 (2014).

41. M. Schäfer, C. Brütting, I. T. Baldwin, M. Kallenbach, High-throughput quantification of more than 100 primary- and secondary-metabolites, and phytohormones by a single solid-phase extraction based sample preparation with analysis by UHPLC–HESI–MS/MS. Plant Methods 12, 30 (2016).

42. Teutloff, Erik, “Balance in plant bacterial recruitment affects resilience to fungal invasion of the phyllosphere,” Friedrich Schiller University of Jena, Jena, Germany. (2023).

43. S. Chen, Y. Zhou, Y. Chen, J. Gu, fastp: an ultra-fast all-in-one FASTQ preprocessor. Bioinformatics 34, i884–i890 (2018).

44. D. Kim, J. M. Paggi, C. Park, C. Bennett, S. L. Salzberg, Graph-based genome alignment and genotyping with HISAT2 and HISAT-genotype. Nat. Biotechnol. 37, 907–915 (2019).

45. H. Li, et al., The Sequence Alignment/Map format and SAMtools. Bioinformatics 25, 2078–2079 (2009).

46. Y. Liao, G. K. Smyth, W. Shi, featureCounts: an efficient general purpose program for assigning sequence reads to genomic features. Bioinformatics 30, 923–930 (2014).

