## Supplementary Material for "Immunity and bacterial recruitment in plant leaves are parallel processes that together shape sensitivity to temperature stress"

**This file includes:**
Supplementary Fig 1-10
Supplementary Table 1

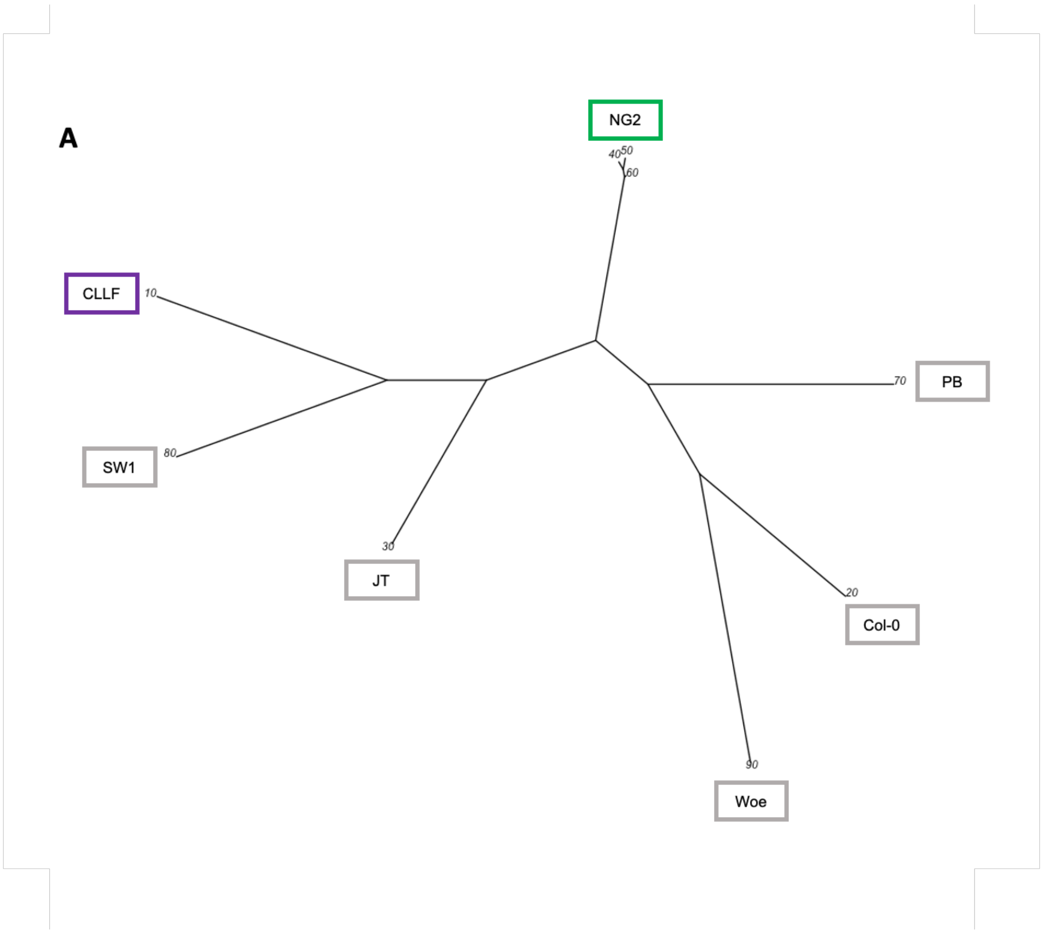


**Supplementary Fig.1. Phylogenetic analysis using whole genome Single Nucleotide Polymorphisms (SNPs) showing unique genotypes of CLLF and NG2.** CLLF and NG2 are labelled using purple and green colors respectively. 10- CLLF, 40,50, and 60 - individual NG2 genomes, 20 – Col-0, 30, 70, 80 and 90 – other locally collected *A. thaliana* ecotypes.

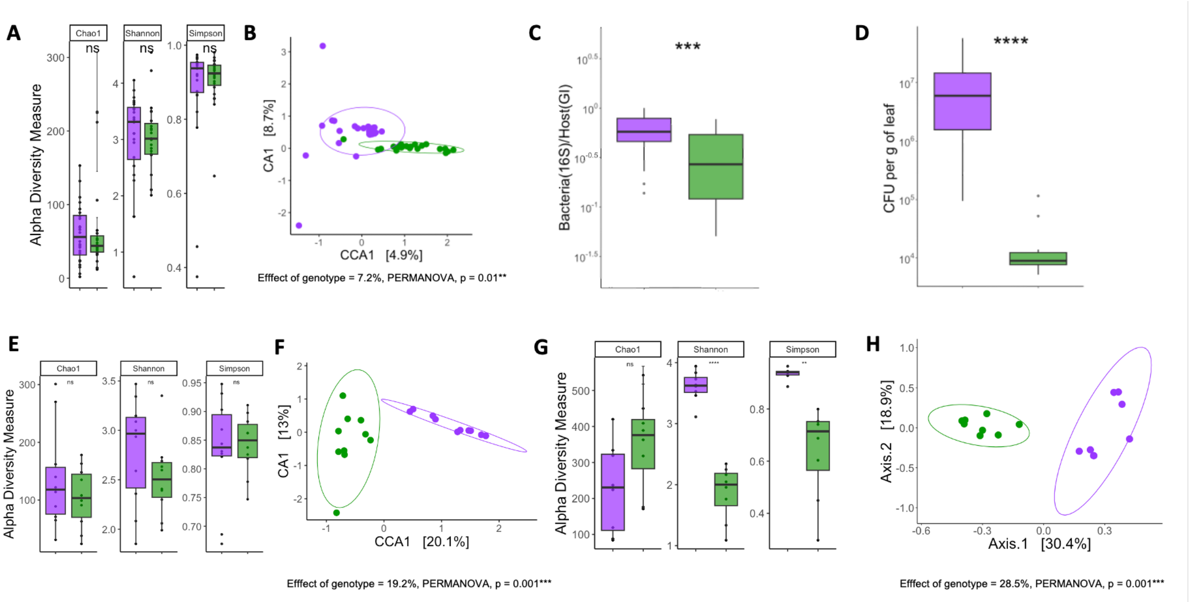


**Supplementary Fig.2. Results from three independent experiments showing altered microbiota recruitment in CLLF.** (A. to D.): <u>Experiment 1.</u> A. Alpha diversity measures of CLLF and NG2 leaf bacteriome based on 16S rRNA gene amplicon sequencing data. B. CCA plot based on the Bray-Curtis distance between samples constrained for genotype, generated using 16S rRNA gene amplicon sequencing data grouped at the genus level. 7.2% variance is explained by genotype (PERMANOVA with 999 permutations) p-value < 0.01. C. Bacteria to host ratio of CLLF and NG2 calculated using hamPCR amplicon sequencing data, the fraction of the total number of bacterial 16S rRNA gene reads to the sum of bacterial 16S rRNA gene and *A. thaliana* GI (*GIGANTEA*) reads per sample (n=24) D. Bacterial loads of CLLF and NG2 Wt measured as colony-forming units (CFU) per gram fresh weight of leaf (NG2, n=12 and CLLF, n=20). (E. and F.): <u>Experiment 2.</u> E. Alpha diversity measures of CLLF and NG2 Wt leaf bacteriome based on 16S rRNA gene amplicon sequencing data. F. CCA plot based on the Bray-Curtis distance between samples constrained for genotype, generated using 16S rRNA gene amplicon sequencing data grouped at the genus level. 19.2% variance is explained by genotype (PERMANOVA with 999 permutations) p-value < 0.001. (G. and H.): <u>Experiment 3.</u> G. Alpha diversity measures of CLLF and NG2 leaf bacteriome based on 16S rRNA gene amplicon sequencing data. H. Principal coordinate analysis (PCoA) based on the Bray-Curtis distance between samples, generated using 16S rRNA gene amplicon sequencing data grouped at the genus level. 28.5% variance is explained by genotype (PERMANOVA with 999 permutations) p-value < 0.001. All amplicon sequencing data are clustered at the OTU level. (CLLF; purple, NG2; green, ns; not significant, * p<0.05, ** p<0.01, *** p<0.001).

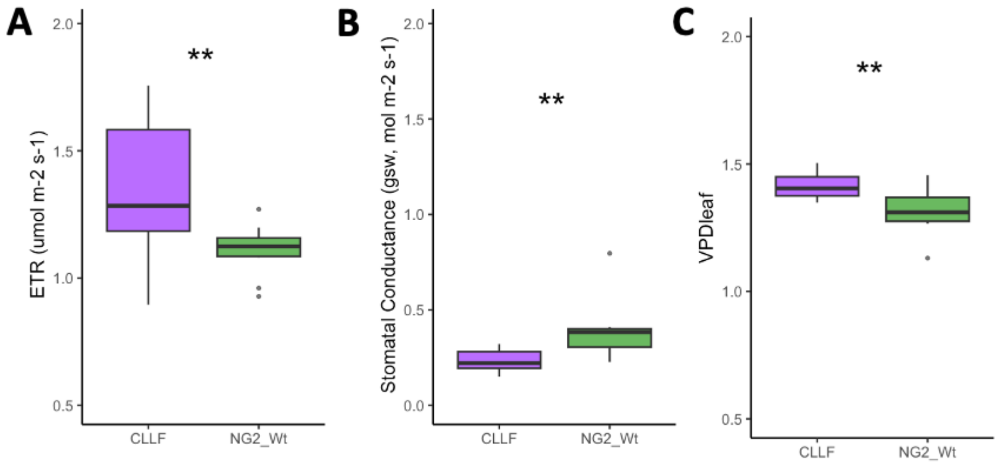


**Supplementary Fig.3. Leaf function measurements of 3-week-old CLLF and NG2 plants.** A. Electron transport rate (ETR). B. Stomatal conductance (gsw). C. Vapor pressure difference between the air and the leaf intercellular air space ($VPD_{leaf}$). (CLLF; purple, NG2 green, ns; not significant, * p<0.05, ** p<0.01, *** p<0.001)

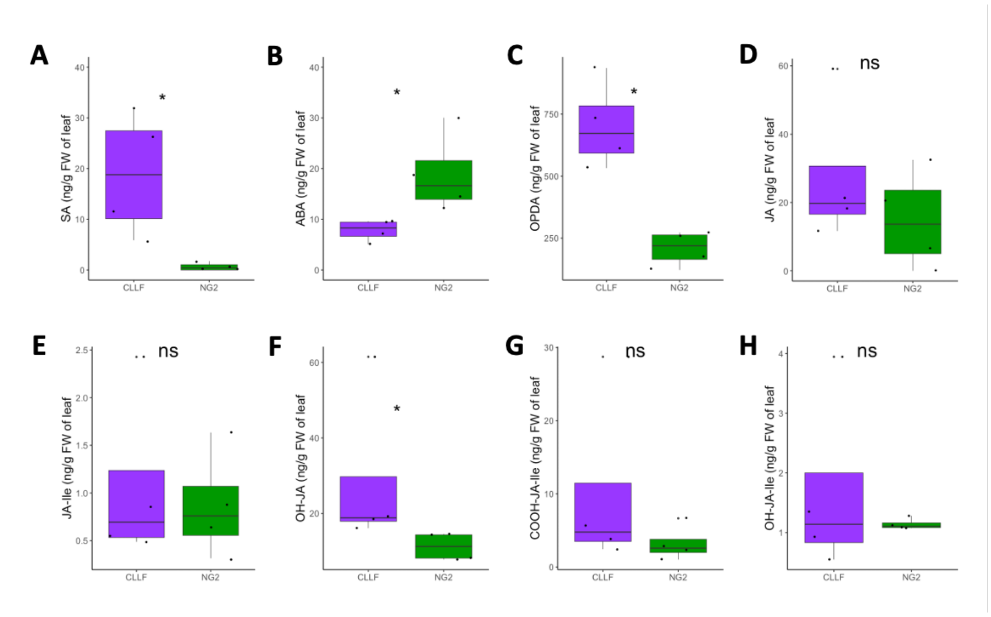


**Supplementary Fig. 4. Phytohormone profiles of CLLF and NG2 plants grown in axenic conditions.** A. Salicylic acid (SA). B. Abscisic acid (ABA). C. Oxylipin 12-oxo-phytodienoic acid (OPDA). D. Jasmonic acid (JA). E. Jasmonic acid-isoleucine (JA-Ile). F. 12-*hydroxy jasmonic acid* (OH-JA). G.12-carboxyl jasmonic acid isoleucine (COOH-JA-Ile). H. 12-*hydroxy jasmonic acid isoleucine (OH-JA-Ile).* All data were measured as nanograms per g fresh weight (n=4, *, $p < 0.05$, ns; not significant)

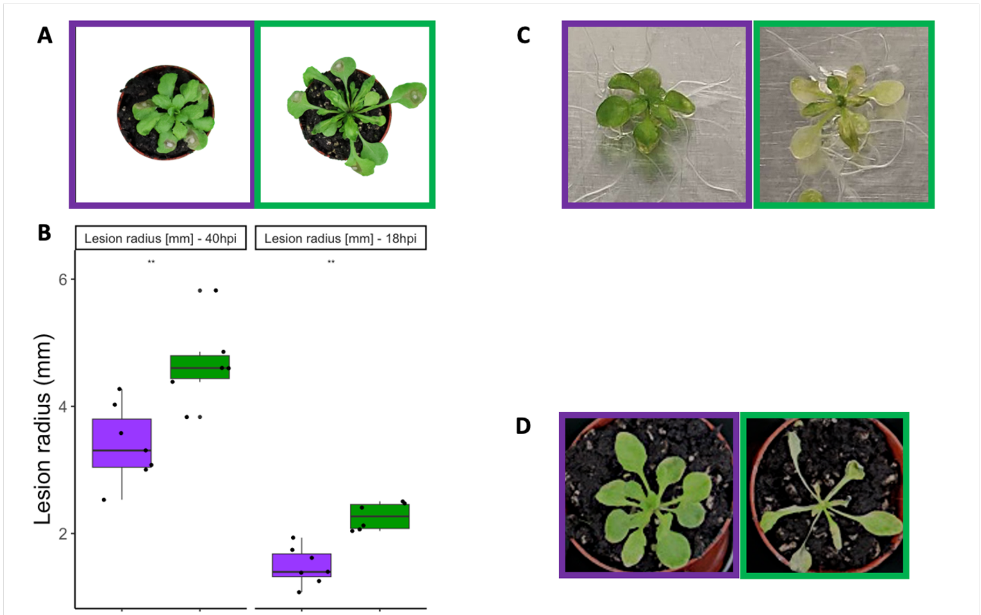


**Supplementary Fig.5.** A. Representative images showing *Sclerotinia sclerotiorum* infection 40 hour-post-infection (hpi). B. Lesion radius of *Sclerotinia sclerotiorum* infected leaves of CLLF and NG2 plants at 40hpi and 18hpi (n=8). C. Representative images of CLLF and NG2 plants flood inoculated with *Pseudomonas syringae pv. syringae* DC3000 48hpi. D. Representative images of CLLF and NG2 plants after extreme heat stress of 34ºC.

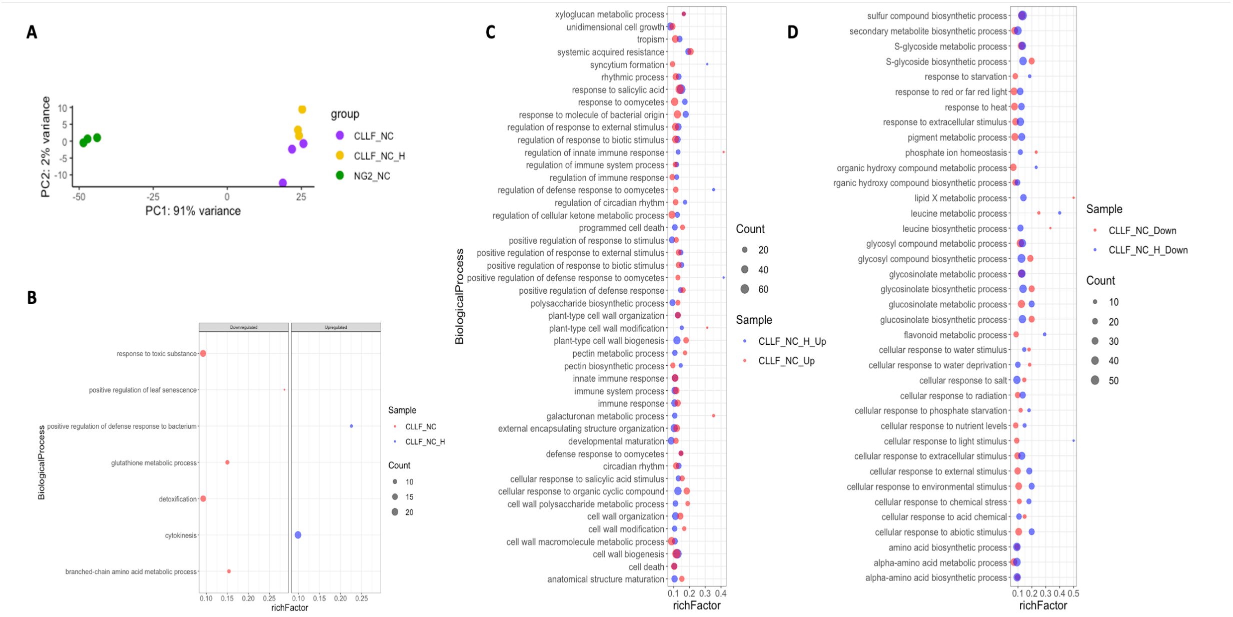


**Supplementary Fig. 6. Gene expression of CLLF plants that before and after heat stress relative to NG2 at normal temperature shows an active immune system.** A. Principal components analysis (PCA) showing variance between differentially expressed genes of 3-week-old CLLF, NG2 plants and CLLF plants that underwent heat stress (30ºC for 48hrs). B. Unique upregulated and downregulated biological processes of CLLF plants with and without heat stress in comparison to NG2 plants. C. Commonly upregulated biological processes in CLLF plants with and without heat stress in comparison to NG2 plants. D. Commonly downregulated biological processes in CLLF plants with and without heat stress in comparison to NG2 plants.

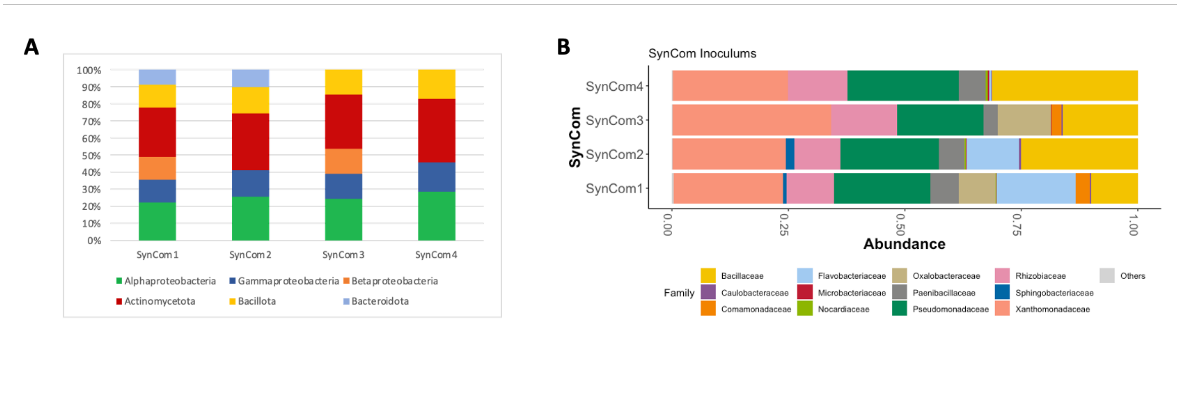


**Supplementary Fig.7. Expected and actual SynCom composition.** A. Bar plot showing an expected composition of the 4 SynComs at the class level. B. Actual composition of SynComs at the family level by 16S rRNA sequencing of SynCom inoculums grown in R2A agar plates.

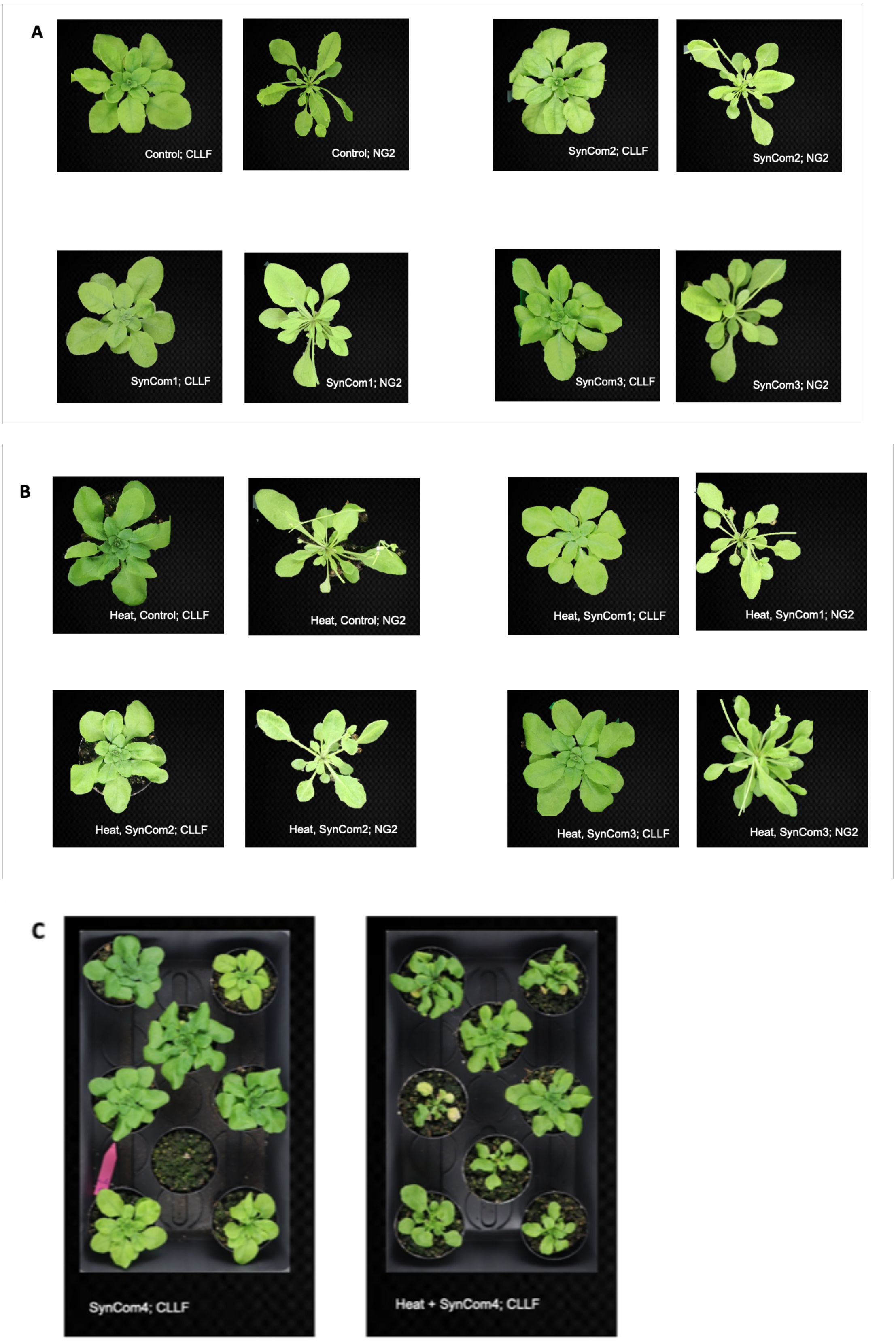


**Supplementary Fig. 8. Representative images of NG2 and CLLF plants showing effects of SynCom and heat stress treatments.** A. Representative CLLF and NG2 plants without a Syncom (control) or with SynCom 1, 2, or 3 in normal conditions. B. Representative CLLF and NG2 plants without a Syncom (control) or with SynCom 1,2 or 3, after heat stress. C. Images of CLLF plants showing effects of heat stress; Left: plants inoculated with SynCom4 without heat stress. Right: plants inoculated with SynCom4 after heat stress.

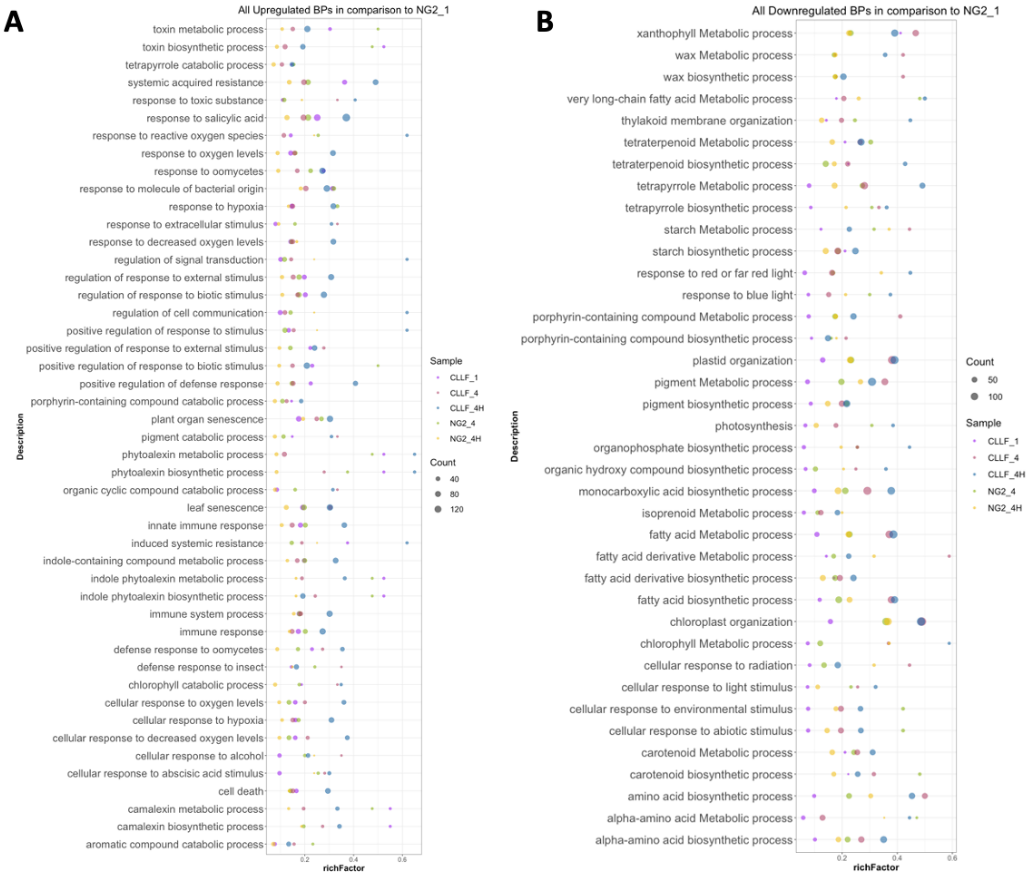


**Supplementary Fig. 9. All common differentially expressing biological processes.** (A. and B.) All up- (A.) or downregulated (B.) biological processes (BPs) in CLLF plants with SynCom1 (Diverse community) at normal temperature (Purple), NG2 and CLLF plants inoculated with SynCom4 at normal temperature (light green and pink, respectively), and NG2 and CLLF plants inoculated with SynCom4 with heat stress (yellow and blue, respectively), all in comparison to NG2 inoculated with SynCom1 at normal temperatures. All genes considered for assigning biological processes are subsetted based on p </= 0.01, basemean > 100, and foldchange > [2]. richFactor is the ratio of differentially expressed genes annotated to the term to the ratio of background genes annotated to the term.

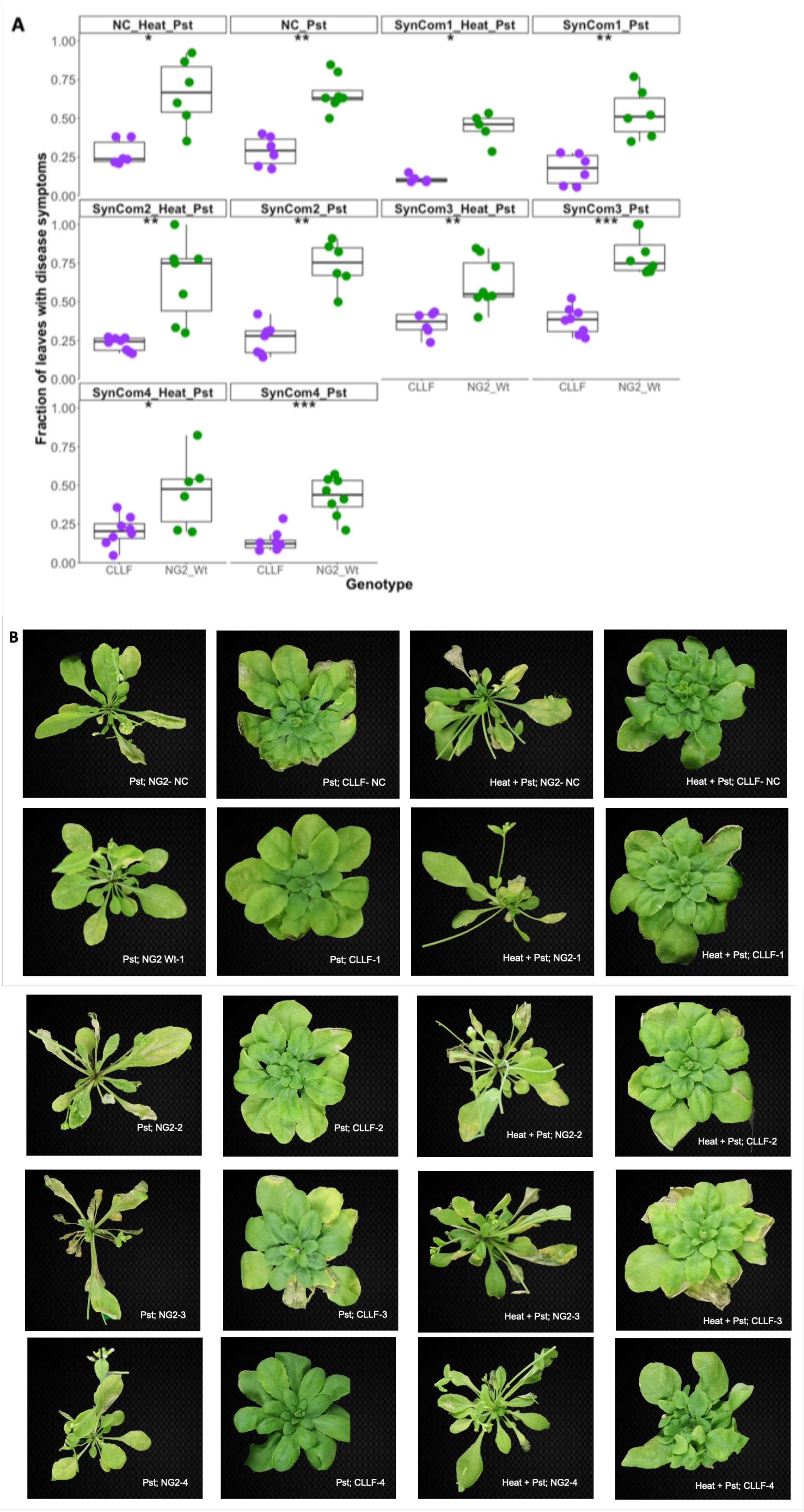


**Supplementary Fig. 10. Effect of SynComs and heat stress on *Pseudomonas syringae pv syringae* DC3000 infection of CLLF and NG2 plants.** A) Fraction of leaves with disease symptoms. B) Representative images of NG2 and CLLF plants infected with *Pseudomonas syringae pv syringae* DC3000. (CLLF; purple, NG2; green, ns; not significant, * p<0.05, ** p<0.01, *** p<0.001).

**Supplementary Table 1. List of SynCom strains used in this study.** All bacterial strains belonging to Bacteroides are colored in blue and all bacteria belonging to Betaproteobacteria are colored in orange. SynCom1 consists of all the 45 strains listed here, SynCom2 lacked Betaproteobacteria, SynCom3 lacked Bacteroides and SynCom4 lacked both Betaproteobacteria and Bacteroides.

| No. | Genus | Species | Family | Class | Phylum | Strain Name |
|---|---|---|---|---|---|---|
| 1 | Nocardia | sp. | Nocardiaceae | Actinomycetia | Actinomycetota | 8_D9 |
| 2 | Plantibacter | flavus | Microbacteriaceae | Actinomycetia | Actinomycetota | SynCom C |
| 3 | Nocardioides | Sp. | Nocardioidaceae | Actinomycetia | Actinomycetota | 3_3 |
| 4 | Microbacterium | foliorum | Microbacteriaceae | Actinomycetia | Actinomycetota | M30_1_10 |
| 5 | Microbacterium | *phyllosphaerae* | Microbacteriaceae | Actinomycetia | Actinomycetota | SynCom E |
| 6 | Curtobacterium | flaccumfaciens | Microbacteriaceae | Actinomycetia | Actinomycetota | 4 E1 |
| 7 | Brevibacterium | frigoritolerans | Brevibacteriaceae | Actinomycetia | Actinomycetota | C19 |
| 8 | Rhodococcus | *degradans* | Nocardiaceae | Actinomycetia | Actinomycetota | SynCom D |
| 9 | Aeromicrobium | sp. | Nocardioidaceae | Actinomycetia | Actinomycetota | 6_4 |
| 10 | Plantibacter | auratus | Microbacteriaceae | Actinomycetia | Actinomycetota | 4 D7 |
| 11 | Agrococcus | jejuensis | Microbacteriaceae | Actinomycetia | Actinomycetota | M30_1_18 |
| 12 | Arthrobacter | agilis | Micrococcaceae | Actinomycetia | Actinomycetota | 5 D2 |
| 13 | Clavibacter | michiganensis | Microbacteriaceae | Actinomycetia | Actinomycetota | 1 E10 |
| 14 | Paenibacillus | amylolyticus | Paenibacillaceae | Bacilli | Bacillota | SynCom F |
| 15 | Bacillus | *thuringiensis* | Bacillaceae | Bacilli | Bacillota | C21 |
| 16 | Bacillus | *aerius* | Bacillaceae | Bacilli | Bacillota | C17 |
| 17 | Bacillus | *nealsonii* | Bacillaceae | Bacilli | Bacillota | C18 |
| 18 | Bacillus | *pumilus* | Bacillaceae | Bacilli | Bacillota | C14 |
| 19 | Bacillus | simplex | Bacillaceae | Bacilli | Bacillota | M30_2_2 |
| 20 | Flavobacterium | *sp.* | Flavobacteriaceae | Flavobacteriia | Bacteroidota | 19_8 |
| 21 | Chryseobacterium | aquaticum | Weeksellaceae | Flavobacteriia | Bacteroidota | C32 |
| 22 | *Chryseobacterium* | polytrichastri | Weeksellaceae | Flavobacteriia | Bacteroidota | 11_B4 |
| 23 | Pedobacter | psychrodurus | Sphingobacteriaceae | Sphingobacteriia | Bacteroidota | C31 |
| 24 | Methylobacterium | phyllostachyos | Methylobacteriaceae | Alphaproteobacteria | Pseudomonadota | 1 C4-1 |
| 25 | Methylorubrum | extorquens | Methylobacteriaceae | Alphaproteobacteria | Pseudomonadota | C36 |
| 26 | Sphingomonas | faeni | Sphingomonadaceae | Alphaproteobacteria | Pseudomonadota | 1 C10-1 |
| 27 | Sphingobium | czechense | Sphingomonadaceae | Alphaproteobacteria | Pseudomonadota | C33 |
| 28 | Rhizobium | sp. | Rhizobiaceae | Alphaproteobacteria | Pseudomonadota | 16_7 |
| 29 | Brevundimonas | intermedia | Caulobacteraceae | Alphaproteobacteria | Pseudomonadota | M30_1_14 |
| 30 | Rhizobium | nepotum | Rhizobiaceae | Alphaproteobacteria | Pseudomonadota | M30_1_5 |
| 31 | Rhizobium | gei | Rhizobiaceae | Alphaproteobacteria | Pseudomonadota | 7 F7 |
| 32 | Agrobacterium | tumefaciens | Rhizobiaceae | Alphaproteobacteria | Pseudomonadota | SynCom B |
| 33 | Bosea | sp. | Boseaceae | Alphaproteobacteria | Pseudomonadota | 6_9 |
| 34 | Rhizobacter | fulvus | incertae sedis | Betaproteobacteria | Pseudomonadota | 2_F6 |
| 35 | Acidovorax | facilis | Comamonadaceae | Betaproteobacteria | Pseudomonadota | M30_4_16 |
| 36 | Variovorax | sp. | Comamonadaceae | Betaproteobacteria | Pseudomonadota | 43_3 |
| 37 | Massilia | arvi | Oxalobacteraceae | Betaproteobacteria | Pseudomonadota | 1 F9 |
| 38 | Janthinobacterium | lividum | Oxalobacteraceae | Betaproteobacteria | Pseudomonadota | A1 |
| 39 | Janthinobacterium | lividum | Oxalobacteraceae | Betaproteobacteria | Pseudomonadota | A2 |
| 40 | Stenotrophomonas | *rhizophila* | Xanthomonadaceae | Gammaproteobacteria | Pseudomonadota | SynCom G |
| 41 | Pseudoxanthomonas | japonensis | Xanthomonadaceae | Gammaproteobacteria | Pseudomonadota | C35 |
| 42 | Pseudomonas | koreensis | Pseudomonadaceae | Gammaproteobacteria | Pseudomonadota | SynCom H |
| 43 | Xanthomonas | campestris/arboricola/hortorum/dyei/cynarae/gardneri | Xanthomonadaceae | Gammaproteobacteria | Pseudomonadota | M30_4_9 |
| 44 | Xanthomonas | campestris | Xanthomonadaceae | Gammaproteobacteria | Pseudomonadota | 1 D3 |
| 45 | Pseudomonas | syringae | Pseudomonadaceae | Gammaproteobacteria | Pseudomonadota | 8 A3 |
